# MuSiC2: cell type deconvolution for multi-condition bulk RNA-seq data

**DOI:** 10.1101/2022.05.08.491077

**Authors:** Jiaxin Fan, Yafei Lyu, Qihuang Zhang, Xuran Wang, Mingyao Li, Rui Xiao

## Abstract

Cell type composition of intact bulk tissues can vary across samples. Deciphering cell type composition and its changes during disease progression is an important step towards understanding disease pathogenesis. To infer cell type composition, existing cell type deconvolution methods for bulk RNA-seq data often require matched single-cell RNA-seq (scRNA-seq) data, generated from samples with similar clinical conditions, as reference. However, due to the difficulty of obtaining scRNA-seq data in diseased samples, only limited scRNA-seq data in matched disease conditions are available. Using scRNA-seq reference to deconvolve bulk RNA-seq data from samples with different disease conditions may lead to biased estimation of cell type proportions. To overcome this limitation, we propose an iterative estimation procedure, MuSiC2, which is an extension of MuSiC [1], to perform deconvolution analysis of bulk RNA-seq data generated from samples with multiple clinical conditions where at least one condition is different from that of the scRNA-seq reference. Extensive benchmark evaluations indicated that MuSiC2 improved the accuracy of cell type proportion estimates of bulk RNA-seq samples under different conditions as compared to the traditional MuSiC [1] deconvolution. MuSiC2 was applied to two bulk RNA-seq datasets for deconvolution analysis, including one from human pancreatic islets and the other from human retina. We show that MuSiC2 improves current deconvolution methods and provides more accurate cell type proportion estimates when the bulk and single-cell reference differ in clinical conditions. We believe the condition-specific cell type composition estimates from MuSiC2 will facilitate downstream analysis and help identify cellular targets of human diseases.

## INTRODUCTION

Bulk tissue RNA sequencing (RNA-seq) technology has been widely adopted to investigate gene expression variations that underlie phenotypic differences among individuals. Findings from these studies have facilitated the identification of therapeutic targets for human diseases. Bulk RNA-seq measures the average gene expression across cells in a tissue and ignores the possible cell-to-cell heterogeneity in the tissue. It is well known that variations not only exist in gene expression patterns across different cell types, but also in cell type compositions across different samples. Knowledge in cell type composition and its changes during disease progression is an important step towards understanding disease pathogenesis [2-4]. Since certain cell types are more vulnerable to diseases than others [5,6], it is important to account for cell type composition variations in bulk RNA-seq data analysis to avoid biasing downstream results [7].

A number of deconvolution methods have been developed to estimate cell type composition in bulk tissue using single-cell RNA-seq (scRNA-seq) data as reference [1,8,9]. A key assumption of these methods is that the bulk samples and scRNA-seq reference samples are from the same population so that the cell-type-specific gene expression observed in the scRNA-seq data can be used to approximate the unobserved cell-type-specific gene expression in the bulk samples. This assumption is reasonable when the bulk and scRNA-seq samples are collected from individuals of similar clinical condition. However, when the bulk samples are collected from individuals with different conditions as compared to the scRNA-seq samples, this assumption becomes questionable. Most existing scRNA-seq data were generated from tissues of healthy donors. When using such data as reference to deconvolve bulk RNA-seq samples collected from diseased subjects [1,3,10], the estimated cell type compositions might not be accurate due to potential differences in cell-type-specific gene expression between healthy and diseased samples and could consequently bias downstream analyses. To better integrate bulk RNA-seq and scRNA-seq data, improved deconvolution methods that are robust to the choice of scRNA-seq reference are needed.

To this end, we extended MuSiC [1] to MuSiC2, which performs deconvolution analysis of bulk RNA-seq data using a scRNA-seq reference data generated from samples with a clinical condition that differs from the bulk data. The key idea is to remove genes from the single-cell reference data that show cell-type-specific differential expression (DE) between different clinical conditions to improve deconvolution and achieve more accurate cell type proportion estimation.

Through extensive benchmark evaluations, we demonstrate that MuSiC2 improved cell type proportion estimation of multi-condition bulk RNA-seq samples as compared to MuSiC [1], which ignored the difference in disease conditions between the bulk and single-cell reference data. We applied MuSiC2 to pancreatic islets bulk RNA-seq data and correctly recovered the trend of cell type proportion changes as type 2 diabetes progressed. We also applied MuSiC2 to human retina bulk RNA-seq data and successfully detected cell type proportion changes with age-related macular degeneration (AMD) progression. With the wide application of large-scale bulk tissue RNA-seq in biomedical studies under different disease conditions, MuSiC2 allows the utilization of existing bulk RNA-seq data to elucidate cell type contributions and identify cellular targets of human diseases.

## METHODS

MuSiC2 is an iterative algorithm that aims to improve cell type deconvolution for bulk RNA-seq data using scRNA-seq data as reference when the bulk data are generated from samples with multiple clinical conditions where at least one condition is different from the scRNA-seq reference. MuSiC2 takes two datasets as input, a scRNA-seq data generated from one clinical condition, and a bulk RNA-seq dataset collected from samples with multiple conditions in which one or more is different from the single-cell reference data. Without loss of generality, we assume the scRNA-seq data are collected from healthy individuals, while the bulk RNA-seq data are collected from healthy and diseased individuals. The key idea of MuSiC2 is that, when the bulk samples and single-cell reference samples are from different clinical conditions, the majority of genes shall still share similar cell-type-specific gene expression pattern regardless of clinical conditions. We refer those genes as “stable” genes, whose cell-type-specific gene expression observed in the scRNA-seq data can be used to approximate the unobserved cell-type-specific gene expression in the bulk samples and to infer the cell type compositions. By removing genes with cell-type-specific differential expression (DE) between samples with different clinical conditions from the single-cell reference, the assumption for cell type deconvolution is met, and hence holds the potential to yield more accurate cell type proportion estimates.

MuSiC2 starts by using MuSiC [1] to infer the subject-specific cell type proportions of the bulk tissue referencing on the scRNA-seq data, and deconvolve the bulk-level expression over the estimated cell type proportions through non-negative least square regression to obtain the cell-type-specific mean expression for each gene. Genes with cell-type-specific DE are detected by comparing the cell-type-specific mean expression between healthy and diseased samples. MuSiC2 then iterates over 2 steps. In Step 1, we use MuSiC [1] to infer the subject-specific estimated cell type proportions of the bulk tissue for healthy samples referencing on the original scRNA-seq data, and for diseased samples referencing on the scRNA-seq data with the DE genes removed. In Step 2, for samples with each condition, we deconvolve the bulk-level expression over the cell type proportion estimates obtained in Step 1 to infer the cell-type-specific mean expression for each gene and detect and remove cell-type-specific DE genes between conditions. By alternating between cell type deconvolution (Step 1) and cell-type-specific DE gene detection and removal (Step 2), MuSiC2 gradually refines the list of “stable” genes retained in the scRNA-seq reference data to improve the cell type proportion estimation for the diseased samples. An overview of MuSiC2 is shown in **Figure 1**.

**Figure 1.**
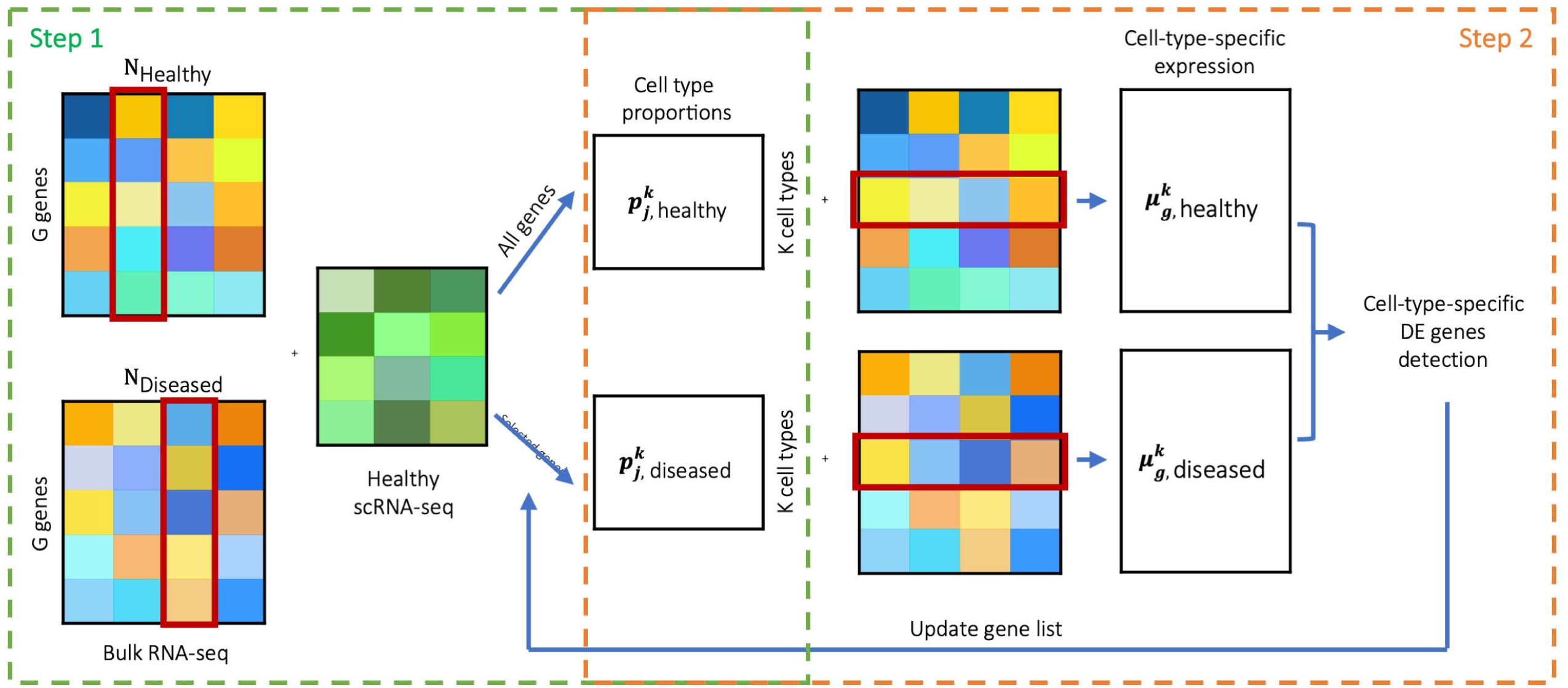
Overview of MuSiC2. The method takes a bulk RNA-seq dataset that includes samples from two different clinical conditions, e.g., healthy and diseased, and a scRNA-seq dataset that includes samples from a single condition, e.g., healthy, as input. MuSiC2 iterates over Steps 1 and 2 and achieves accurate cell type proportion estimation after the algorithm converges. In Step 1, under each condition, we employ MuSiC [1] to infer the cell type proportions, 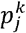, of the bulk tissue for individual *j* of cell type *k* by borrowing information from scRNA-seq reference data from healthy individuals. When deconvolving healthy bulk samples, all genes in the single-cell reference are used, whereas only “stable” genes are used for diseased samples. At the initial step, all genes are included for cell type deconvolution in Step 1. In Step 2, for samples of each condition (e.g., healthy and diseased), we model bulk-level expression of gene g across individuals over the estimated cell type composition from Step 1, 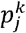, through non-negative least square regression to infer the cell-type-specific mean expression, 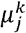. By comparing 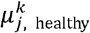 to 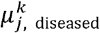, we then identify genes with cell-type-specific DE between the two conditions, and update the gene list in the scRNA-seq reference data used in Step 1 by removing those cell-type-specific DE genes for the diseased sample deconvolution.

### Step 1: Estimating cell type proportions by deconvolution

In this step, we infer cell type proportions of bulk RNA-seq samples of each condition by incorporating cell-type-specific gene expression information provided by the scRNA-seq reference data collected from healthy samples. Following the notation in MuSiC [1], for samples of each condition, the bulk-level total read count of gene *g* for individual *j,X*_*jg*_, can be written as a weighted sum of *K* cell-type-specific gene expressions,

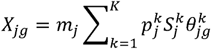

where, for individual *j,m*_*j*_ is the total number of cells in the tissue for bulk RNA-seq, 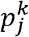 is the proportion of cells from cell type 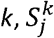 is the average cell size of cell type *k*, and 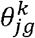 is the relative abundance of gene *g* for cell type *k*. Borrowing information from the scRNA-seq reference, subject-level 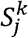 and 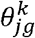 can be approximated by the sample average *S*^*k*^ and 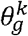. By modeling the total expression across *G* genes, *X*_*jg*_ (*g* = 1, … *G*,), over *S*^*k*^ and 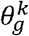, the subject-specific cell type proportion,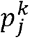, can be estimated through weighted non-negative least squares regression, denoted by 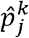.

For bulk RNA-seq data of healthy samples, as the scRNA-seq data are also generated from healthy subjects, we use all genes in the single-cell reference for deconvolution. While for diseased samples, due to potential discrepancy in cell-type-specific gene expression profiles between the bulk and scRNA-seq samples, we only use those “stable” genes in the single-cell reference for deconvolution. We define a “stable” gene as a gene that has similar expression profiles across all cell types between the bulk and scRNA-seq samples regardless of the clinical conditions. The key of deconvolution for diseased bulk samples lies in the selection of those “stable” genes.

### Step 2: Detecting cell-type-specific DE genes

In this step, under each condition, by using the cell type proportion estimates obtained in Step 1, for each gene, we infer its cell-type specific mean expression and determine if this gene shows evidence of differential expression in any given cell type. Under each condition, the bulk-level total read counts for gene *g, X*_*jg*, healthy_ and *X*_*jg*, diseased_, can be written as weighted sums of *K* cell-type-specific gene expressions:

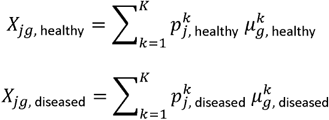

where 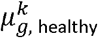 and 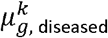 are the mean expression of gene for cell type for healthy and diseased samples, respectively. For healthy samples, as 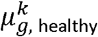 should be ≥0 for a given gene *g*, it can be inferred through a non-negative least squares regression by modeling the bulk-level expression across all healthy individuals, *X*_*jg*, healthy_ (*j* = 1, …, *N*_healthy_) on the estimated cell type proportions from Step 1, 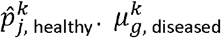 can be inferred similarly using diseased samples.

For a given gene *g* in cell type *k*, we use the log fold change of cell-type-specific expression between conditions, 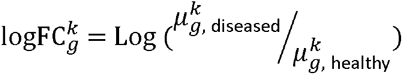, to test for the cell-type-specific DE by employing a resampling procedure in order to achieve a reliable estimate. Specifically, at each iteration, we generate a subset of samples by random sampling without replacement under each condition with same sampling proportion in each condition, and compute the 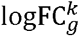. We define a statistic 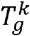 as the ratio of the mean and standard deviation (SD) of the 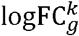 over all resampling iterations as a measure of the cell-type-specific DE. Genes with 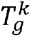 in the top 5% for common cell types, i.e., cell types with average proportion ≥ 10%, or in the top 1% for rare cell types, i.e., cell types with average proportion < 10%, are considered as cell-type-specific DE genes. Since fold change is sensitive to genes with low expression, we suggest that genes with bulk-level average sequencing depth < 20 are retained as ‘stable’ genes and excluded from the cell-type-specific DE detection. We further filter the genes by their expression levels in the random samples. Specifically, we compute the mean of 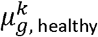 and 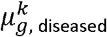 over the resampling iterations, and retain genes with cell-type-specific expression in the bottom 5% for samples in both conditions as ‘stable’ genes and exclude them from the cell-type-specific DE detection.

We update the scRNA-seq data used for deconvolution of the diseased samples in Step 1 by removing the cell-type-specific DE genes from the single-cell reference. Iterating through the deconvolution and DE gene detection steps, MuSiC2 gradually refines the “stable” gene set and eventually achieves more accurate cell type proportion estimates after the algorithm converges. For individual *j*, we denote the cell type proportion estimates of cell type *k* as 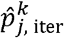 for the current iteration and 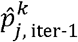 for the previous iteration. A cell type *k* is said to converge when 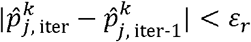 for a rare cell type and 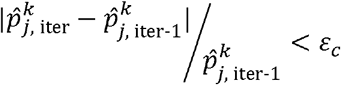 for a common cell type, where *ε*_*r*_ and *ε*_*c*_ are two pre-specified convergency cutoffs used for the rare and common cell types, respectively. An individual sample is considered to converge when all *K* cell types have converged. MuSiC2 will stop the iteration and be considered to converge when all but ≤ 1 samples of the bulk RNA-seq data have converged.

### Benchmark bulk RNA-seq data generation

To evaluate the performance of MuSiC2, we constructed benchmark bulk RNA-seq data in which the proportion of each cell type is known, using a publicly available scRNA-seq dataset on human pancreatic islets generated from six healthy individuals and four type 2 diabetes (T2D) patients [11]. We evaluated the performance of MuSiC2 as a function of sample size and cell type proportions. For each condition, i.e., healthy and T2D, the number of individuals was set to be 20, 50 or 100. For each individual in the benchmark bulk RNA-seq data, we generated 500 single cells from six selected cell types, including acinar, alpha, beta, delta, ductal and gamma, with cell type proportions generated from a Dirichlet distribution, where the mean parameters were informed from an existing data to mimic what are seen in real world. Specifically, we obtained the condition-specific cell type proportions in the Fadista *et al*. bulk RNA-seq samples [12] using the single-cell data with matched condition [11] as reference. The cell type proportion estimates were then used as the mean parameters of the Dirichlet distribution to generate the cell type proportions for healthy and diseased samples separately (**Figure 2A**).

**Figure 2.**
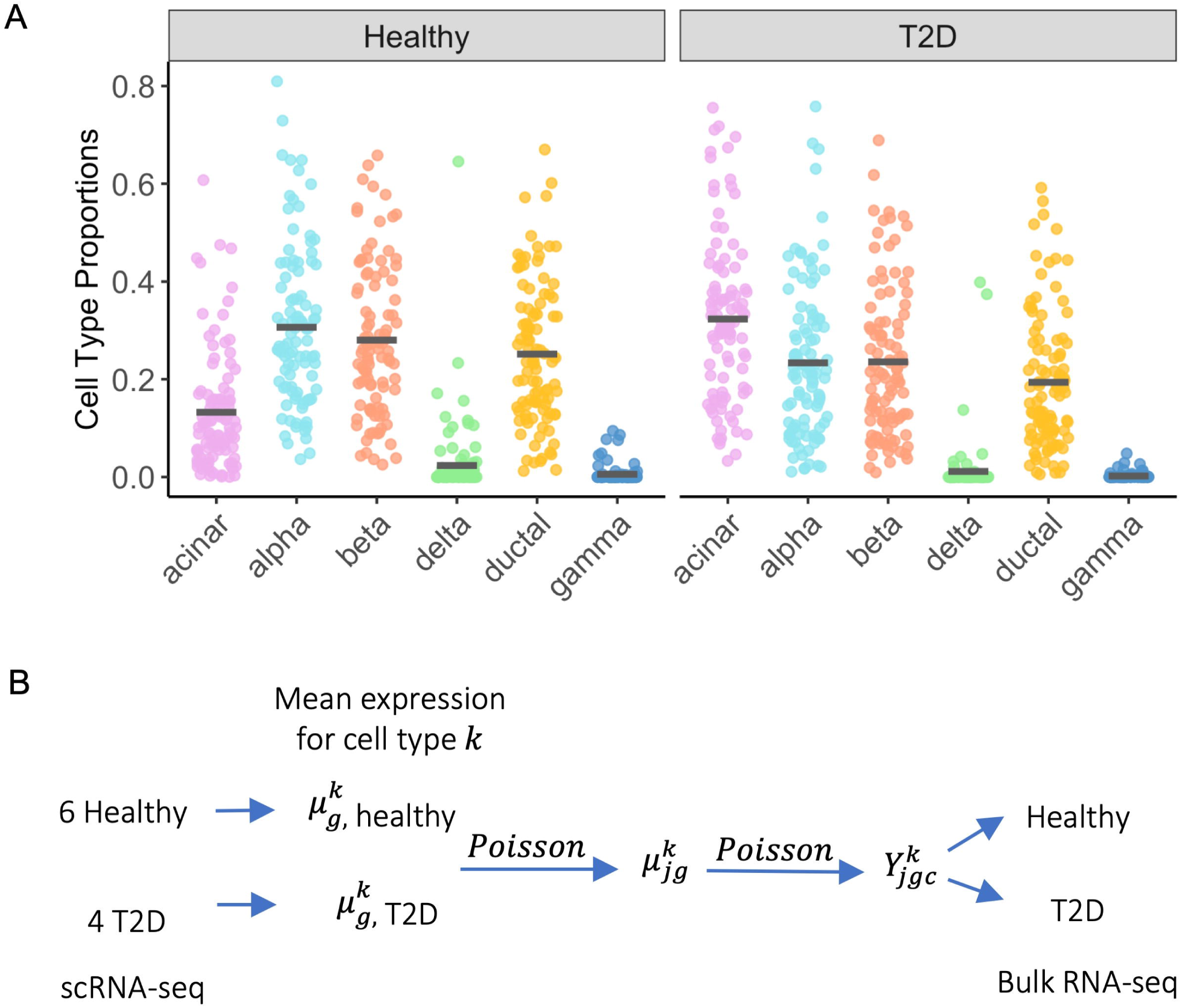
Benchmark evaluation data generation scheme. **(A)** Cell type compositions for the artificial bulk RNA-seq data for healthy and T2D samples. For each cell type, the solid line represents the average cell type proportions across samples, which was inferred through bulk data deconvolution. **(B)** Data generation scheme for obtaining the artificial bulk RNA-seq data. Given gene *g*, for each cell type *k*, we first inferred the condition-specific mean expression level from the single-cell data [11]. We then sampled the subject-specific mean expression through Poisson distribution with the inferred mean as the rate parameter. Another layer of Poisson sampling was employed to obtain the total read count of cell *c* for samples of each condition by using subject-specific mean as the rate. The artificial bulk RNA-seq data was obtained by summing up the read counts across all cells.

For each cell, we simulated read counts for 10,000 genes randomly selected from genes that were expressed in at least 30 cells in the scRNA-seq data [11]. **Figure 2B** shows the overall workflow for benchmark data generation. Briefly, to simulate data for healthy samples, for each gene, we obtained the average cell-type-specific gene expression across cells from 6 healthy donors in the scRNA-seq data [11]. The subject-specific mean expression was sampled from a Poisson distribution with mean set to the inferred cell-type-specific mean expression from the single-cell data [11]. The cell-specific total read counts for each individual were then sampled from another Poisson distribution with the mean determined by the subject-specific mean expression. By summing up total read counts across all cells, we obtained the benchmark bulk-level gene expression data. Following the same scheme, we simulated benchmark bulk RNA-seq data for diseased individuals using the scRNA-seq data from the 4 T2D donors [11].

### Evaluation metrics

For individual *j*, denote the true cell type proportions as *p*_*j*_ and estimated cell type proportions as 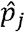, where *p*_*j*_ and 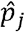 are vectors with length *K*. We evaluated the deconvolution methods using the following metrics:

i. 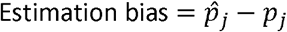
ii. Root mean squared deviation, 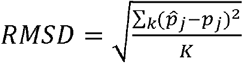

## RESULTS

### Benchmark evaluation

We performed benchmark studies to evaluate the performance of MuSiC2. The benchmark bulk RNA-seq data included healthy and diseased (i.e., T2D) samples generated from a human pancreatic islet scRNA-seq dataset [11], with known cell type proportions under each condition. We evaluated the performance of MuSiC2 as a function of sample size, which took values of 20, 50 and 100 per condition. As a comparison, we also performed the deconvolution analysis using MuSiC [1]. Specifically, we compared four scenarios: deconvolving healthy and diseased bulk samples first using healthy and then using the diseased scRNA-seq data as reference. The estimated cell type proportions were compared to the true proportions to evaluate the accuracy of the deconvolution methods.

With 50 individuals per condition, when deconvolving healthy samples using healthy scRNA-seq data as reference, MuSiC and MuSiC2 had exactly the same performance as expected with estimation bias close to 0, because MuSiC2 only refines the gene list in the single cell reference when the bulk and single cell data are from different conditions. However, as diseases often progress gradually, even healthy individuals can be heterogeneous in terms of the potential towards developing diseases. Therefore, we considered removing the cell-type-specific DE genes from the single cell reference and updating the cell type proportions through iterations when bulk and single cell reference data are both from healthy individuals. However, the results did not improve much. This is likely due to the fact that some of the detected cell-type-specific DE genes are cell-type-specific marker genes and removal of those marker genes leads to less accurate estimates in the cell type proportion for healthy bulk samples. Therefore, we recommend to update the reference genes and cell type proportions using MuSiC2 only if the bulk and single cell reference samples are collected from different disease conditions. In contrast, when deconvolving healthy samples with diseased single cell reference, MuSiC generated biased proportion estimates especially for acinar, beta and ductal cells. We observed a severe overestimation for beta cells and underestimation for ductal cells. Whereas MuSiC2 significantly improved the proportion estimation for the three cell types by removing the DE genes in the single cell reference. Especially for beta cells, MuSiC2 even outperformed the deconvolution results obtained by MuSiC with healthy single cell reference, highlighting the significance of gene selection for deconvolution (**Figure 3A**).

**Figure 3.**
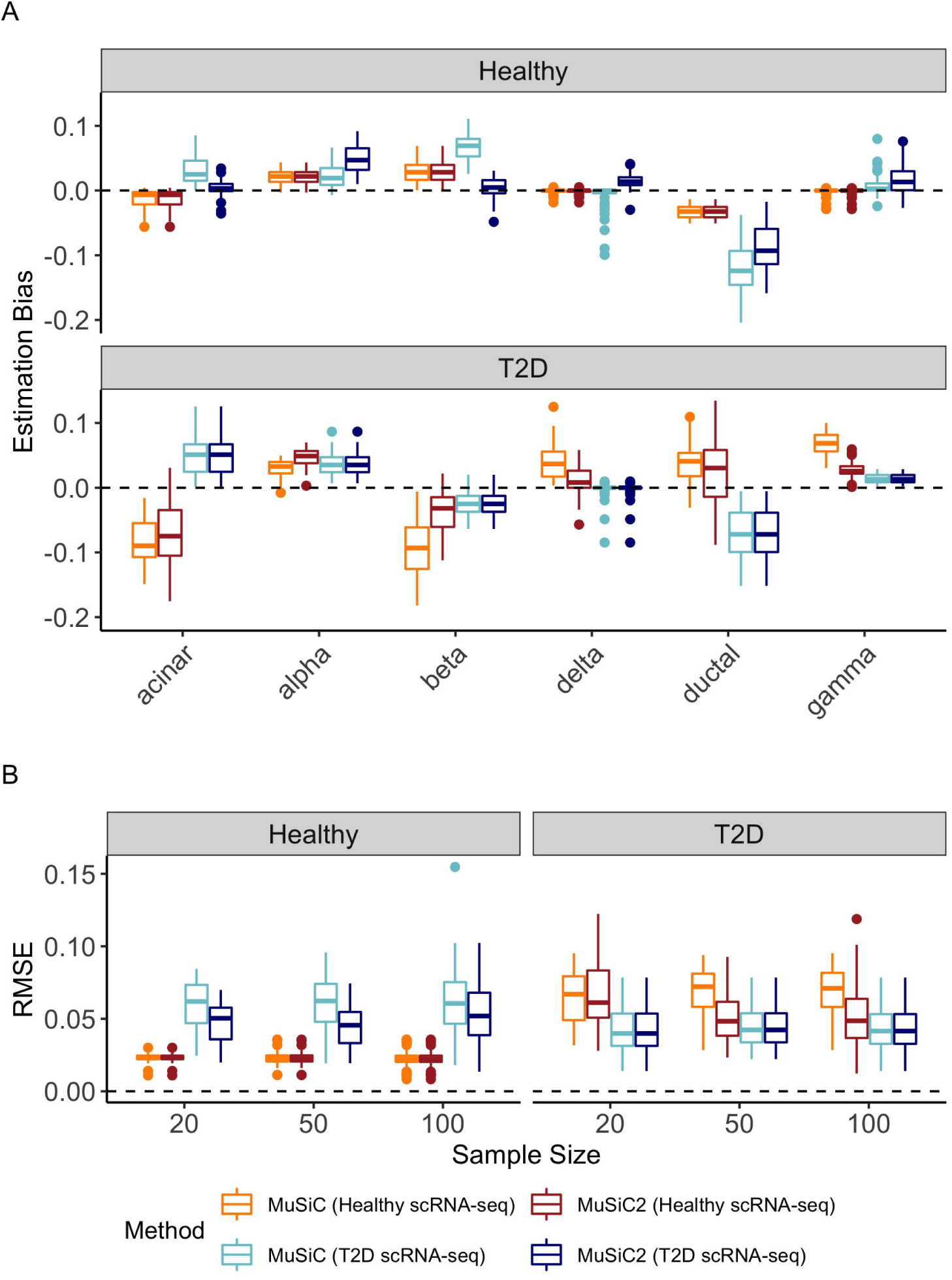
Estimation accuracy of benchmark datasets. We evaluated the performance of MuSiC2 and compared to MuSiC using the bulk RNA-seq data of healthy and T2D sample, with four scenarios were considered: MuSiC and MuSiC2 with healthy scRNA-seq reference, and MuSiC and MuSiC2 with diseased scRNA-seq reference. **(A)** Cell-type level estimation bias of the four scenarios to deconvolve healthy and T2D bulk samples, with 50 individuals in each condition. **(B)** Boxplot of individual-level root mean square error (RMSE) of the four scenarios to deconvolve healthy and T2D bulk samples as a function of sample size per condition.

Similarly, when deconvolving diseased samples using diseased scRNA-seq reference, the performance was the same between MuSiC2 and MuSiC. However, the estimation bias is larger as compared to deconvolving healthy individuals using healthy scRNA-seq reference, which could potentially be due to the larger variability in diseased samples. In contrast, when deconvolving diseased samples with healthy single-cell reference, compared to MuSiC, MuSiC2 improved the estimation accuracy for most of the cell types. Particularly for beta cells, a common cell type in islets, MuSiC2 produced much more accurate cell type proportion estimates than MuSiC, which suffered from severe underestimation.

Interestingly, for alpha and ductal cells, when deconvolving the diseased samples, the performance of MuSiC with diseased scRNA-seq reference was worse than the situation when using healthy scRNA-seq reference. We suspect this is likely due to a larger variation in cell-type-specific expression for alpha and ductal cells in the diseased single-cell reference as compared to the healthy single cell reference (**Figure 3A**). Similar patterns were observed when sample size per condition set to 20 and 100 (**Supplementary Figure S1**).

In addition to the cell-type-level estimation bias, we further evaluated the performance of MuSiC2 using individual-level root mean square error (RMSE) across cell types as a function of sample size per condition. For both healthy and diseased individuals, as expected, the estimation accuracy was higher when deconvolving samples using a single cell reference with matched clinical condition as compared to using a scRNA-seq from a different condition. Specifically, when bulk samples and single cell reference were from different conditions, compared to MuSiC, MuSiC2, on average, reduced the RMSE by over 20% under most scenarios investigated. As sample size varied, not surprisingly, we noticed that the performance of MuSiC was consistent regardless of the sample size as MuSiC deconvolved each individual separately. Meanwhile, even though with some variability, MuSiC2, in general, had a comparable performance in terms of the RMSE for deconvolving healthy bulk samples using diseased scRNA-seq data regardless of the sample size per condition. In contrast, when deconvolving diseased bulk samples with healthy scRNA-seq data, as diseased samples were more variable than healthy samples, the performance of MuSiC2 improved when sample size increased from 20 to 50. However, when sample size further increased from 50 to 100, the performance of MuSiC2 did not improve much (**Figure 3B**). In real studies, we often have less diseased samples than healthy individuals. Thus, we also considered a scenario assuming 30 T2D versus 70 healthy individuals, and observed similar patterns of estimation bias and RMSE (**Supplemental Figure S2**).

Next, we evaluated the replicability of MuSiC2 by applying MuSiC2 to the same benchmark bulk RNA-seq samples twice. Taking the data with 50 samples per condition for example, we found that the results between the two runs were highly correlated, with an average Pearson correlation coefficient of 0.985 of the estimated cell type proportions across diseased samples deconvolved with healthy single cell reference, and 0.997 across healthy samples deconvolved with diseased single cell reference. Moreover, we studied the effect of the sample size of resampling on the performance of MuSiC2 as we employed the resampling procedure to achieve a reliable measurement of cell-type-specific DE using the log fold change of cell-type-specific gene expression between conditions. Again taking the benchmark data with sample size of 50 per condition for example, when deconvolving diseased bulk RNA-seq data with healthy scRNA-seq reference, MuSiC2 obtained similar cell type proportion estimates when randomly drawing 25%, 50% or 75% of all samples at each resampling iteration (**Supplemental Figure S3**). In addition, we evaluated the cutoff for cell-type-specific DE genes identification by using more stringent cutoffs of removing top 10% and 5% genes ranked by the log fold change rather than top 5% and 1% for the common and rare cell types, respectively. We found that the estimated cell type proportions with less ‘stable’ genes retained in the single cell reference for deconvolution were highly correlated with the results obtained with more ‘stable’ genes retained, suggesting that MuSiC2 is robust to the choice of cutoffs for removing cell-type-specific DE genes. Especially for common cell types, acinar, alpha, beta and ductal, the Pearson correlations were nearly 1. The correlation for gamma cell was slightly lower with a Pearson correlation of 0.82, which was not surprising as gamma cells were extremely rare in the islets samples (**Supplemental Figure S4**).

### Application to human pancreatic islet bulk RNA-seq data

We applied MuSiC2 to human pancreatic islet bulk RNA-seq data generated in an expression quantitative trait loci (eQTL) study by Fadista *et al*. [12]. We chose to reanalyze this data because previous study deconvolved all both healthy and diseased individuals using scRNA-seq data from healthy donors [1]. As shown by our benchmark evaluations, deconvolving diseased samples using healthy single-cell reference may yield biased proportion estimates and such bias may propagate further in downstream analysis. We focused our analysis on 77 samples with measured hemoglobin A1c (HbA1c), a well-known biomarker for T2D with higher HbA1c level indicating higher risk of developing T2D. We defined an individual as T2D if HbA1c ≥ 5.7. We chose 5.7 rather than the traditional cutoff of 6.5 in order to have enough diseased samples. Actually individuals with HbA1c of 5.7 are generally considered as pre-diabetic and their gene expression profiles may have started differing from healthy subjects [13,14]. With this definition of T2D, we ended up with 39 healthy and 38 T2D samples. Six well-recognized cell types were selected: acinar, alpha, beta, delta, ductal and gamma, and scRNA-seq data generated from six healthy donors by Segerstolpe *et al*. [11] were used as the reference.

With healthy scRNA-seq data as reference, we deconvolved the Fadista bulk RNA-seq data [12] using MuSiC and MuSiC2. As a comparison, we also deconvolved the diseased bulk samples using MuSiC with the T2D scRNA-seq as reference. The estimated cell type proportions of the diseased samples obtained using the three methods were summarized in **Figure 4 and Supplementary Table S1**. Compared to the estimated proportions obtained using MuSiC with T2D single cell reference, MuSiC2 outperformed MuSiC when using healthy single cell data as the reference, especially for beta cells. Similar to what we have observed in benchmark studies, with unmatched single cell reference, MuSiC tended to underestimate the beta cell proportions for diseased samples as compared to MuSiC2. As subjects with higher HbA1c tend to have greater risk of developing T2D, we expect to see a negative association between HbA1c levels and beta cell proportions since beta cells are gradually lost during T2D progression [1,15]. If deconvolving Fadista bulk RNA-seq samples [12] using healthy single cell reference as in the MuSiC paper [1], we would overestimate the negative correlation between HbA1c level and beta cell proportions (Spearman correlation = −0.345, p-value = 0.002), as compared to the correlation of the proportion estimates of obtained by MuSiC referenced on matched single cell data (Spearman correlation = −0.227, p-value = 0.047). When using MuSiC2, even with unmatched single cell reference, we were able to successfully recover a similar negative association between beta cell proportions and HbA1c levels (Spearman correlation = −0.239, p-value = 0.036) (**Figure 5**). In fact, through single-cell transcriptome profiling of islets, Lawlor *et al*. have showed that the observed decrease in beta cell type proportion for T2D was not as significant as previously reported [16].

**Figure 4.**
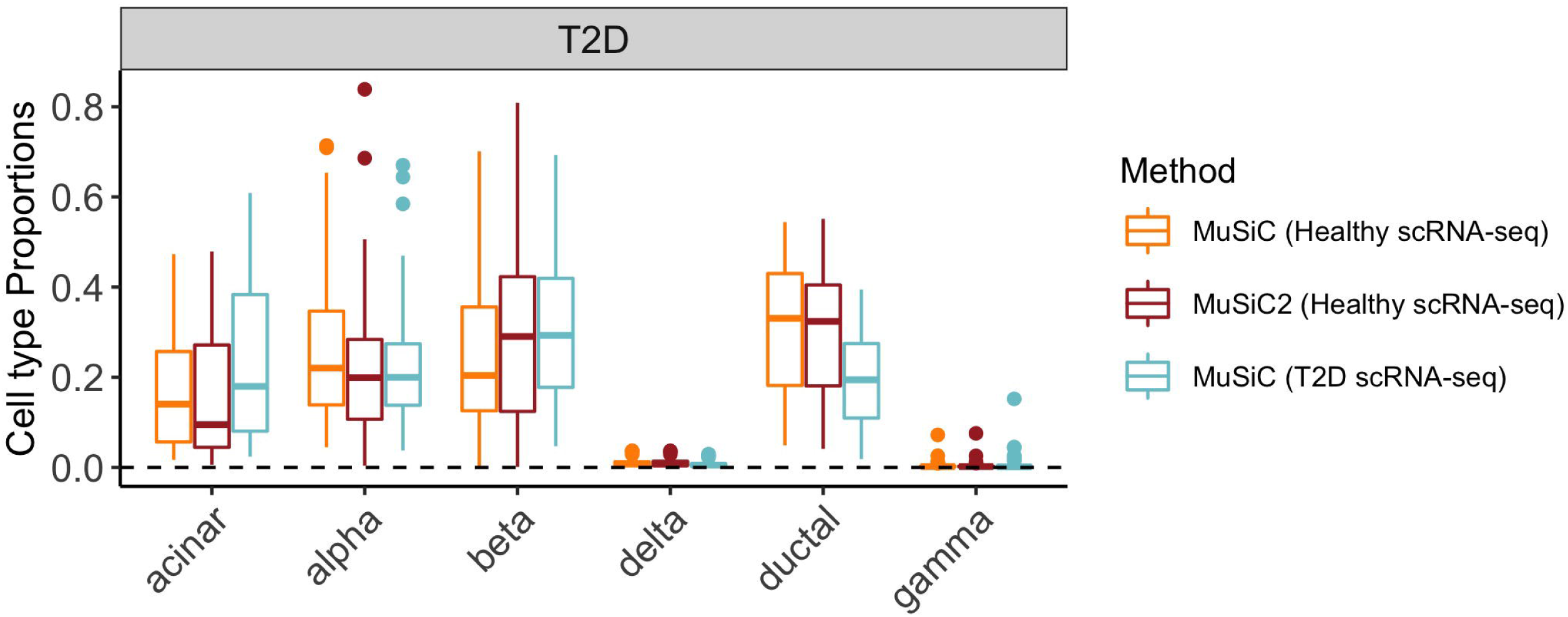
Pancreatic islet cell type composition in diseased samples. Boxplots showing estimated cell type proportions of diseased bulk islets samples [12], using three methods: MuSiC and MuSiC2 with healthy scRNA-seq reference, and MuSiC with diseased scRNA-seq reference.

**Figure 5.**
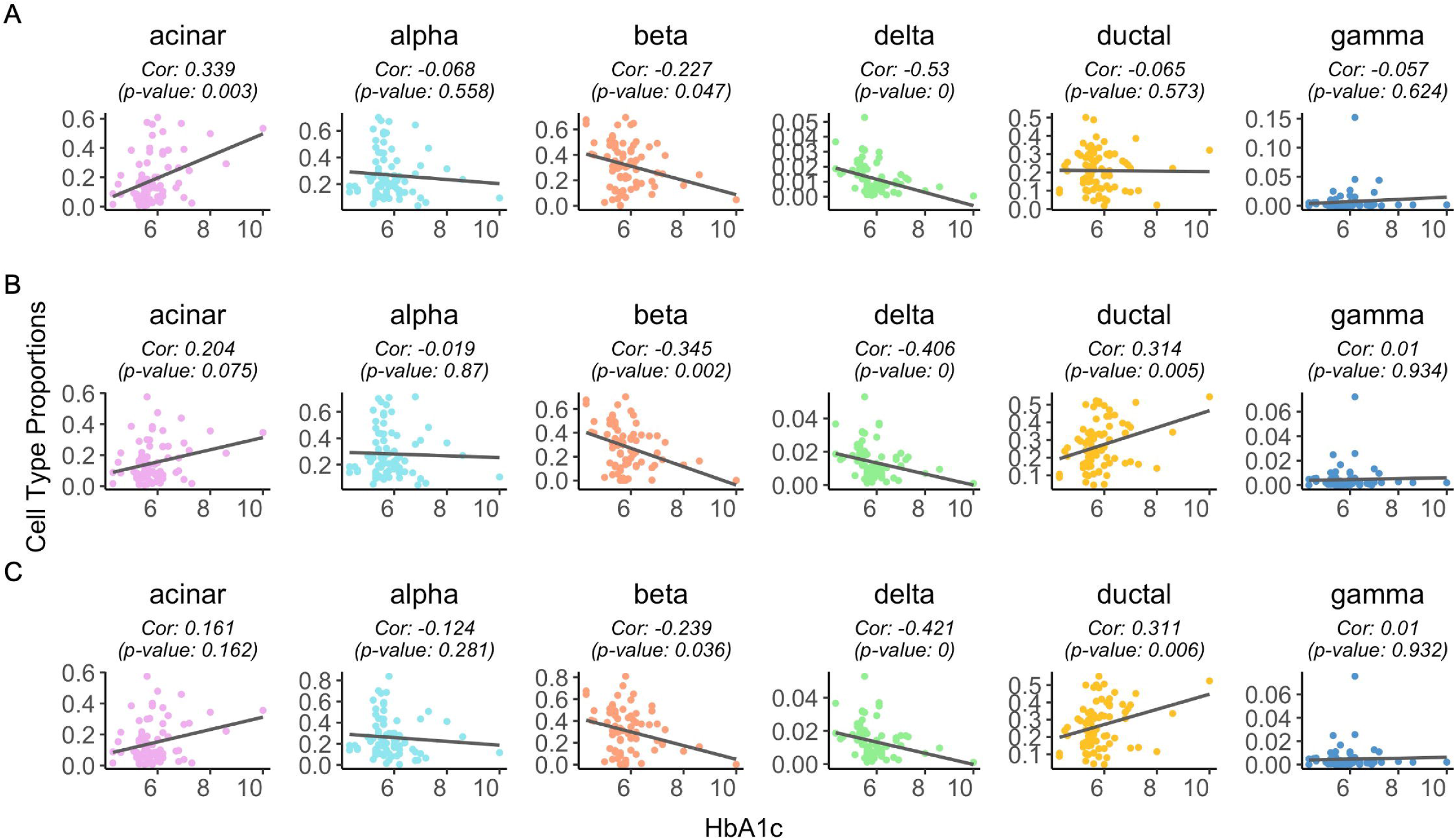
Correlation of HbA1c level and cell type composition of pancreatic islet data. Scatter plot of HbA1c vs. estimated proportions of the pancreatic islet bulk RNA-seq samples [12] for each cell type with a fitted regression line. For each cell type, we calculated the Spearman’s correlation (p-value) between HbA1c and estimated cell type proportions shown as the subtitle of each plot. The proportions were estimated based on three methods: MuSiC with the matched scRNA-seq reference for each condition samples (**A**), MuSiC (**B**) and MuSiC2 (**C**) with healthy scRNA-seq reference.

### Application to human retina bulk RNA-seq data

We further applied MuSiC and MuSiC2 to a human retina RNA-seq data [17]. Age-related macular degeneration (AMD) is a leading cause of blindness and affects over 10 million Americans [18]. Currently with no proven treatments, vision loss is an eventual outcome for many individuals with AMD. To better understand the molecular mechanisms of AMD progression, Lyu *et al*. studied cell type proportional changes in bulk tissues with different AMD stages using deconvolution analysis [19]. However, as single cell RNA-seq data of human retina with specific AMD stages are limited, the single cell data generated from non-AMD donors were used as the reference for deconvolving all bulk samples ignoring the disease status. As unmatched single cell reference may bias the proportion estimates, in this analysis, we aimed to reassess retina cell type composition changes during AMD progression using MuSiC2. Using the human peripheral retina bulk RNA-seq data generated by Ratnapriya *et al*. [20],we performed deconvolution analysis on the selected 105 non-AMD samples and 61 samples carrying moderate-late AMD condition (Minnesota Grading System (MGS) 4)[21]. The single cell RNA-seq data of the same tissue, generated from two non-AMD donors by Lyu *et al*. [19], were used as the reference, which contained cellular gene expression of 6,820 cells across 11 cell types.

**Figure 6** showed cell type composition in non-AMD and AMD peripheral retina, obtained by applying MuSiC [1] and MuSiC2 respectively. We noticed that MuSiC2 better captured the cell type proportion changes in AMD macula. First of all, the result from MuSiC2 revealed a substantial drop of rod cell population in AMD retina (0.303) compared to control (0.481). Such tendency is consistent with the published histological evidence --- the degeneration and loss of rod cells in AMD eyes [22,23]. While the loss of rod cells happens gradually during AMD progression, it is expected to be obvious in MGS3 AMD retina and could be captured by the cell type deconvolution analysis. Additionally, MuSiC2 detected a significant increase (p<0.001) in astrocyte proportion in AMD retina (AMD: 0.109 vs. Control: 0.059). The increase in astrocyte cell population may be associated with the immune response of the retina during AMD progression [24].

**Figure 6.**
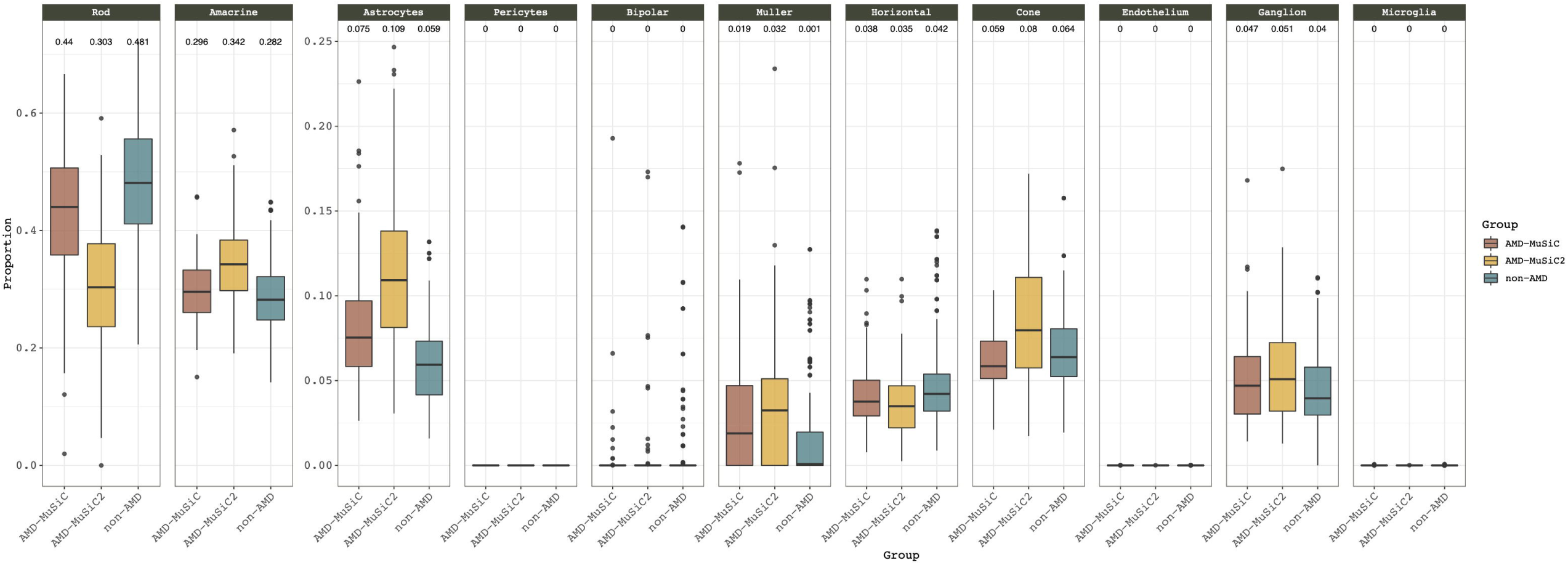
Cell type composition in peripheral retina samples. Boxplots comparing cell type proportions of peripheral retina bulk RNA-seq samples between two different disease status (non-AMD and AMD) [19], estimated using MuSiC and MuSiC2 with non-AMD scRNA-seq reference. The median of cell type proportions across samples is shown as the number above each boxplot.

Compared to MuSiC2, the cell type composition in AMD retina obtained from MuSiC is more akin to the non-AMD retina, especially for the 3 major cell types, inducing rod, amacrine, and astrocytes. This is not surprising as MuSiC ignores potential disease-associated differences in cell-type-specific gene expression when performing deconvolution analysis on AMD retina using scRNA-seq reference from non-AMD retina. This is especially true when major cell types (e.g., rod cell) are vastly impacted by the disease, as an inaccurate estimation of major cell type proportion will affect the estimation of all other minor cell types. In comparison, MuSiC2 took into account the potential discrepancy in gene expression between AMD samples and non-AMD reference by removing cell-type-specific DE genes from the single cell reference, which led to more accurate estimates and better reflected the overall cell type proportion changes with AMD progression.

## DISCUSSION

Insights into cellular heterogeneity and their composition in diseased relevant tissues are crucial for understanding cell-type-specific gene expression variations associated with disease progression and can help identify cellular targets of diseases. Existing cell type deconvolution methods are based on the assumption that bulk and single-cell RNA-seq data are generated from samples under similar clinical conditions and therefore have similar cell-type-specific gene expression profiles. However, only limited scRNA-seq data are available, with the majority of them generated from healthy individuals. Due to the lack of diseased scRNA-seq data, most of the current deconvolution analyses infer the cell type proportions of the bulk RNA-seq data using single-cell reference sequenced from healthy samples, ignoring the difference in disease status between the bulk and single cell reference samples. This could potentially result in biased cell type proportion estimates. To address this issue and to better utilize existing multi-condition bulk RNA-seq data, we proposed MuSiC2, an extension of MuSiC [1], to deconvolve bulk RNA-seq data using single-cell reference generated under a different condition.

MuSiC2 iterates over deconvolution and cell-type-specific DE gene detection. The key idea is that by removing genes with different expression profiles between different clinical conditions from the single-cell reference, we can use the rest of the genes, i.e., ‘stable’ genes, to deconvolve the bulk samples. Over iterations, MuSiC2 refines the list of cell-type-specific DE genes and subsequently improves the cell type proportion estimation. Through extensive benchmark evaluations and application to a pancreatic islet dataset and an eye dataset, we showed that our method improved cell type proportion estimates when the bulk and single-cell reference data were generated from samples with different disease status. One advantage of MuSiC2 is that it does not require large sample size to achieve a reasonably good performance. In addition, it is important to note that, in MuSiC2, the criteria for defining cell-type-specific DE genes are purely data-driven, which offers the users great flexibility of making their own definition of DE genes under their research context. In general, we recommend to only remove the most significant cell-type-specific DE genes for an easier convergence, e.g., the top 5% for common cell types and 1% for rare cell types.

MuSiC2 was evaluated only under scenarios where the bulk and single-cell samples are sequenced from the same species. However, as there are more scRNA-seq data available from animal studies than human studies, the same workflow can be applied to cross-species deconvolution, e.g., deconvolving human bulk RNA-seq data using mouse single-cell reference. In addition, the current implementation of MuSiC2 only handles bulk data with two discrete conditions, e.g., diseased vs. healthy. However, in reality, diseases progress continuously, thus it is possible to have multiple intermediate disease stages or there are no established cutoffs to clearly define disease stages based on quantitative clinical measures. For example, HbA1c measures the glucose level and is used to characterize the risk of type 2 diabetes. It is possible that cell-type-specific gene expression changes along with the HbA1c level as disease progresses. Information may be lost by simply dichotomizing the HbA1c level by a cutoff. In pancreatic islets data analysis example, even though using cutoff of 5.7 separated the samples into healthy and diseased groups with reasonable sizes, however, the resulted diseased group included individuals who were pre-diabetic, and the changes in gene expression profiles from healthy to pre-diabetes may be less as compared to the true T2D individuals defined using cutoff of 6.5. This may therefore dilute the cell-type-specific expression differences between healthy and diseased samples in our analysis. Another example is AMD. In addition to healthy and advanced AMD, there are early and intermediate AMD stages during the macular degeneration. To overcome this limitation, we plan to modify MuSiC2 by identifying genes whose cell-type-specific expression is significantly associated with continuous clinical covariates/stages as measures of disease progression in future studies.

MuSiC2 addresses the gene expression differences between bulk and single cell reference data under two different conditions. However, due to sample heterogeneity, even when the bulk and single cell reference data are generated from similar clinical conditions, there can still be differences in the gene expression profiles between the bulk and single cell reference. Through benchmark studies, we have shown that, with MuSiC2, deconvolving bulk samples using single cell reference from a different clinical condition sometimes can yield more accuracy proportion estimates compared to using single cell reference from similar condition. For future work, we plan to improve the gene selection algorithm for MuSiC2 that takes into account the heterogeneity across samples to further refine the proportion estimates.

In summary, we have developed MuSiC2, an iterative workflow, to perform deconvolution analysis of bulk RNA-seq data using a scRNA-seq reference data generated from samples with a clinical condition that differs from the bulk data. MuSiC2 provides a more accurate cell type proportion estimates when the bulk and single-cell reference differ in clinical conditions compared to MuSiC [1]. Although our method uses MuSiC [1] for cell type proportion estimation in this paper, the general idea of MuSiC2, i.e., removing cell-type-specific DE genes from single cell reference, can be applied to other deconvolution methods. To the best of our knowledge, MuSiC2 is the first deconvolution method that deals with the situation where the bulk and single-cell RNA-seq data are generated from samples with different clinical conditions. We believe this method will facilitate the investigations of different cell types’ contribution to human diseases.

## CODE AVAILABILITY

MuSiC2 is implemented as a R package and is available at https://github.com/Jiaxin-Fan/MuSiC2.

