## Supplemental Figures for "MuSiC2: cell type deconvolution for multi-condition bulk RNA-seq data"

### Supplementary Figures

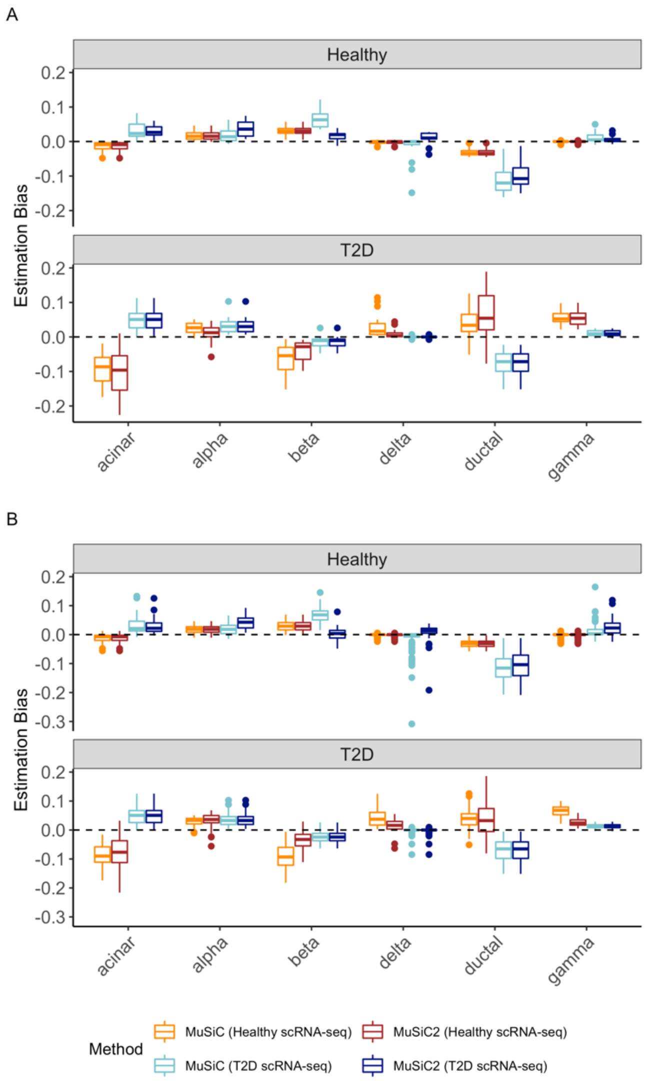


**Figure S1. Estimation accuracy of benchmark datasets with balanced samples.** We evaluated the performance of MuSiC2 and compared it to MuSiC deconvolution method using the bulk RNA-seq data of healthy and T2D samples. In total four deconvolution methods of the multi-condition bulk samples were considered: MuSiC and MuSiC2 with healthy scRNA-seq reference, and MuSiC and MuSiC2 with diseased scRNA-seq reference. Cell-type level estimation bias of the four methods deconvolving healthy and T2D bulk samples were showed in Boxplots, where **(A)** 20 and **(B)** 100 individuals were assumed per condition.

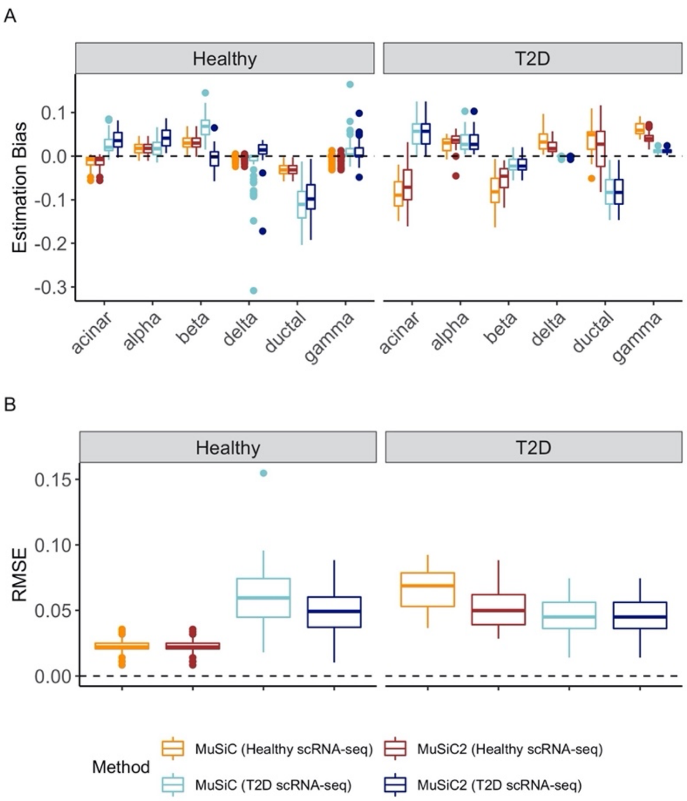


**Figure S2. Estimation accuracy of benchmark datasets with unbalanced samples.** We evaluated the performance of MuSiC2 and compared it to MuSiC deconvolution method using the bulk RNA-seq data containing 70 healthy and 30 T2D individuals. In total four deconvolution methods of the multi-condition bulk samples were considered: MuSiC and MuSiC2 with healthy scRNA-seq reference, and MuSiC and MuSiC2 with diseased scRNA-seq reference. **(A)** Cell-type level estimation bias of the four methods deconvolving healthy and T2D bulk samples. **(B)** Boxplot of individual-level root mean square error (RMSE) of the four methods deconvolving healthy and T2D bulk samples.

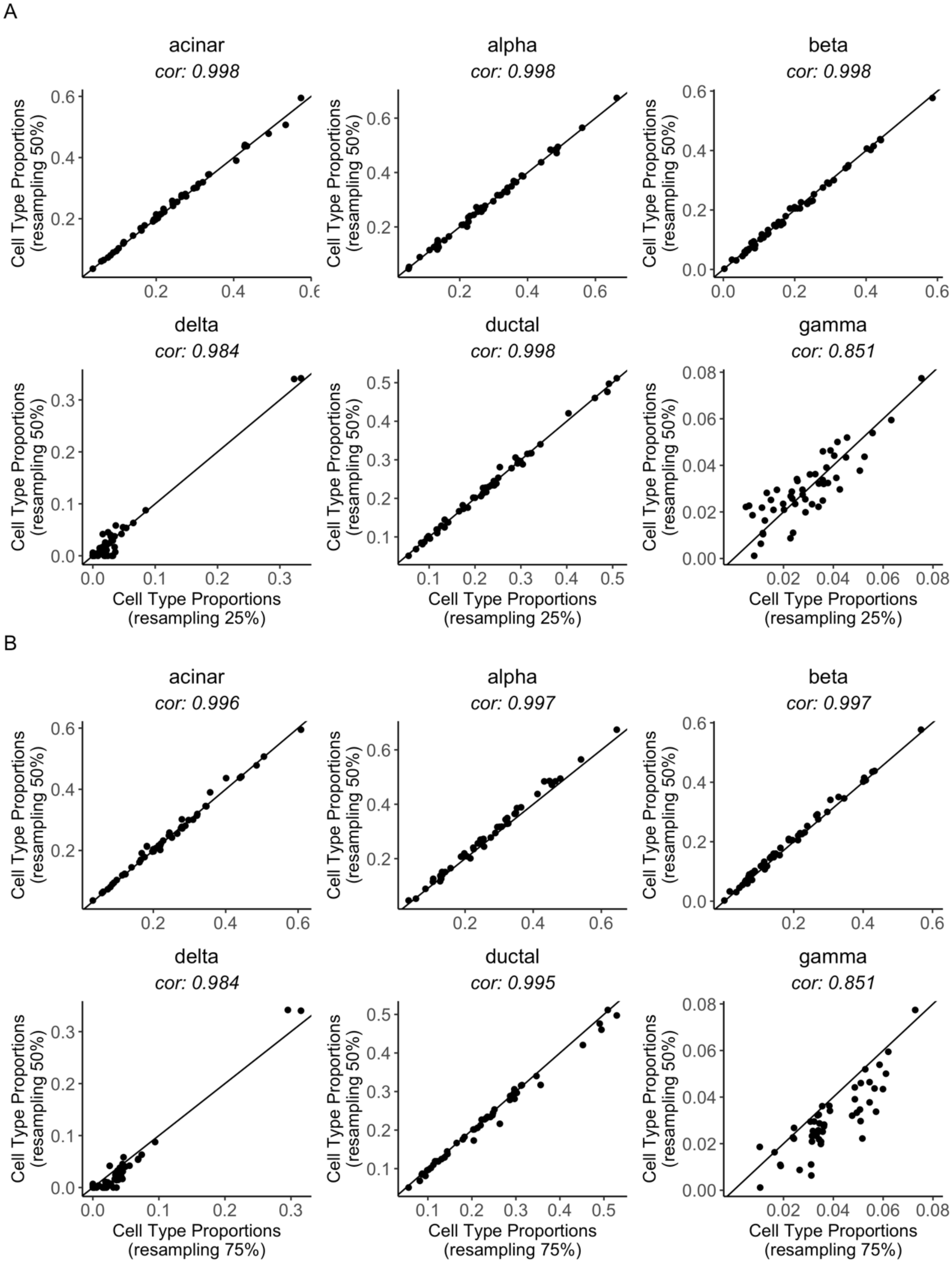


**Figure S3. Evaluation of MuSiC2 regards to sample size of resampling.** We evaluated the consistency of MuSiC2 with respect to the sample size of resampling. The bulk RNA-seq data contained 50 healthy and 50 diseased individuals and were deconvolved with healthy scRNA-seq reference using MuSiC2. For each cell type, scatterplots were made to compare estimated cell type proportions for the 50 diseased individuals obtained by sampling 50% from the original samples, keeping the sampling proportion the same between the two conditions, against proportion estimates obtained by sampling 25% **(A)** or 75% **(B)**. The Pearson correlation coefficients were shown as the subtitle of each plot.

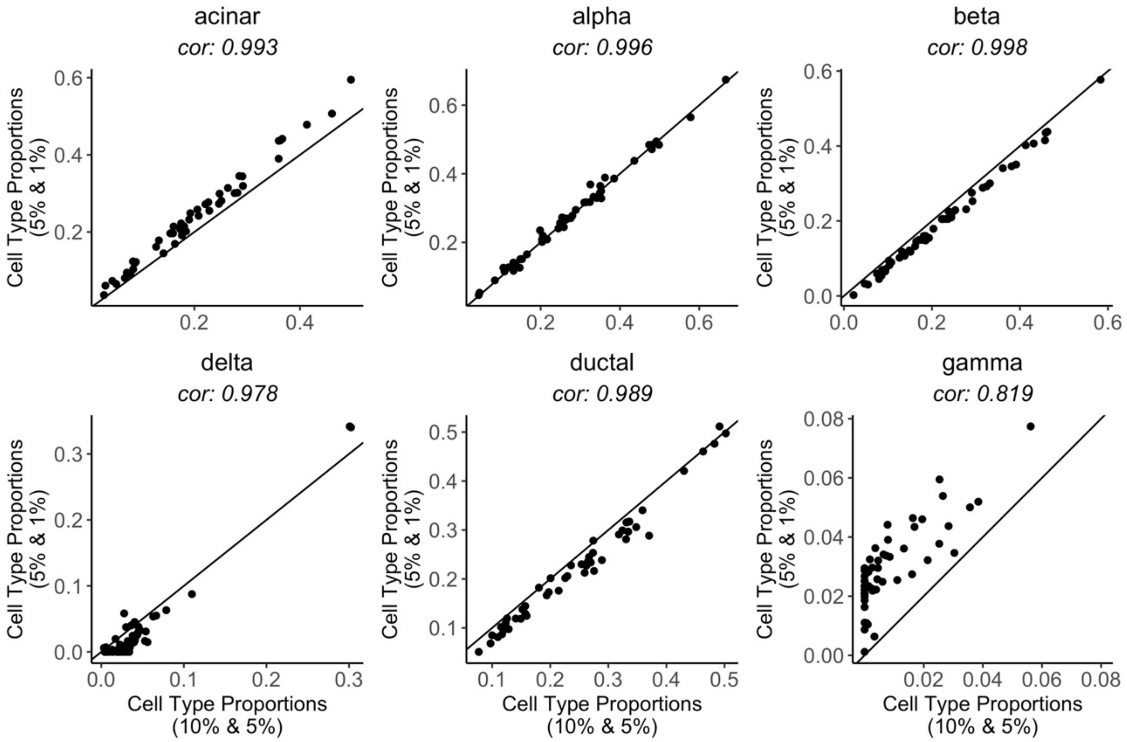


**Figure S4. Evaluation of MuSiC2 regards to cutoffs of cell-type-specific DE genes identification.** We evaluated the consistency of MuSiC2 with respect to the cutoffs used to identify cell-type-specific DE genes. The bulk RNA-seq data contained 50 healthy and 50 diseased individuals and were deconvolved with healthy scRNA-seq reference using MuSiC2. Two sets of cutoffs were considered. One we assumed genes with log fold change statistic $T$ in the top 5% for common cell types or in the top 1% for rare cell types are considered as cell-type-specific DE genes. The other is more stringent, where genes with $T$ in the top 10% for common cell types or in the top 5% for rare cell types are considered as cell-type-specific DE genes. For each cell type, we drew scatterplots comparing estimated cell type proportions obtained using less stringent cutoffs against proportion estimates obtained using more stringent cutoffs, with the Pearson correlation coefficients showed as the subtitle of each plot.
