## Supplemental Table 1 for "MuSiC2: cell type deconvolution for multi-condition bulk RNA-seq data"

**Table S1. Pancreatic islet cell type composition in healthy and diseased samples.** Estimated cell type proportions obtained using three different methods: MuSiC and MuSiC2 with healthy scRNA-seq reference, and MuSiC with the matched scRNA-seq reference for each condition samples.

| Subject | HbA1c | Cell Type | Estimated Cell Type Proportions | | |
| --- | --- | --- | --- | --- | --- |
|  |  |  | MuSiC | MuSiC2 | MuSiC |
|  |  |  | (Matched scRNA-seq data) | (Healthy scRNA-seq) | (Healthy scRNA-seq) |
| Sub1 | 5.8 | acinar | 0.15742009 | 0.133001561 | 0.095976854 |
| Sub1 | 5.8 | alpha | 0.202134978 | 0.297897257 | 0.207916751 |
| Sub1 | 5.8 | beta | 0.302693747 | 0.012640779 | 0.174602969 |
| Sub1 | 5.8 | delta | 0.001332511 | 0.003069877 | 0.002203054 |
| Sub1 | 5.8 | ductal | 0.335333019 | 0.551393034 | 0.517711668 |
| Sub1 | 5.8 | gamma | 0.001085655 | 0.001997493 | 0.001588704 |
| Sub11 | 6.9 | alpha | 0.037593573 | 0.043762926 | 0.045177774 |
| Sub11 | 6.9 | beta | 0.150450799 | 0.131771802 | 0.127481242 |
| Sub11 | 6.9 | acinar | 0.568020312 | 0.458915217 | 0.438441827 |
| Sub11 | 6.9 | delta | 0.002126334 | 0.002566391 | 0.00270627 |
| Sub11 | 6.9 | ductal | 0.241808983 | 0.362983426 | 0.386191786 |
| Sub11 | 6.9 | gamma | 0 | 2.37E-07 | 1.10E-06 |
| Sub13 | 5.4 | ductal | 0.297790292 | 0.297698632 | 0.297790292 |
| Sub13 | 5.4 | delta | 0.017101079 | 0.017105827 | 0.017101079 |
| Sub13 | 5.4 | beta | 0.490542309 | 0.490649931 | 0.490542309 |
| Sub13 | 5.4 | gamma | 0.010911508 | 0.010909381 | 0.010911508 |
| Sub13 | 5.4 | acinar | 0.069425516 | 0.069430115 | 0.069425516 |
| Sub13 | 5.4 | alpha | 0.114229297 | 0.114206113 | 0.114229297 |
| Sub14 | 5.8 | acinar | 0.562255349 | 0.401739509 | 0.383823678 |
| Sub14 | 5.8 | beta | 0.065483638 | 0.001608028 | 0.001586907 |
| Sub14 | 5.8 | delta | 0.000919047 | 0.001682744 | 0.001636398 |
| Sub14 | 5.8 | alpha | 0.09126778 | 0.107589388 | 0.103970794 |
| Sub14 | 5.8 | ductal | 0.278283543 | 0.485073848 | 0.506842647 |
| Sub14 | 5.8 | gamma | 0.001790641 | 0.002306483 | 0.002139576 |
| Sub15 | 5.5 | ductal | 0.185269868 | 0.185044518 | 0.185269868 |
| Sub15 | 5.5 | gamma | 0 | 0 | 0 |
| Sub15 | 5.5 | acinar | 0.063375139 | 0.063375427 | 0.063375139 |
| Sub15 | 5.5 | delta | 0.020310084 | 0.020316348 | 0.020310084 |
| Sub15 | 5.5 | beta | 0.054635505 | 0.054435266 | 0.054635505 |
| Sub15 | 5.5 | alpha | 0.676409404 | 0.676828441 | 0.676409404 |
| Sub16 | 4.6 | acinar | 0.084012327 | 0.084028088 | 0.084012327 |
| Sub16 | 4.6 | alpha | 0.146364064 | 0.146399483 | 0.146364064 |
| Sub16 | 4.6 | beta | 0.494221343 | 0.494111265 | 0.494221343 |
| Sub16 | 4.6 | gamma | 0.002995169 | 0.002995163 | 0.002995169 |
| Sub16 | 4.6 | delta | 0.016601694 | 0.016609373 | 0.016601694 |
| Sub16 | 4.6 | ductal | 0.255805402 | 0.255856627 | 0.255805402 |
| Sub17 | 6.1 | delta | 0.029737092 | 0.035033298 | 0.036077892 |
| Sub17 | 6.1 | beta | 0.646836594 | 0.647282105 | 0.546986626 |
| Sub17 | 6.1 | acinar | 0.029851206 | 0.022827578 | 0.019688073 |
| Sub17 | 6.1 | alpha | 0.163754066 | 0.084661791 | 0.19353457 |
| Sub17 | 6.1 | gamma | 0.003507586 | 0.002390488 | 0.002426168 |
| Sub17 | 6.1 | ductal | 0.126313455 | 0.20780474 | 0.201286671 |
| Sub18 | 5.3 | alpha | 0.131938743 | 0.131955935 | 0.131938743 |
| Sub18 | 5.3 | beta | 0.538094393 | 0.538056141 | 0.538094393 |
| Sub18 | 5.3 | delta | 0.017812767 | 0.017819941 | 0.017812767 |
| Sub18 | 5.3 | gamma | 0.002585754 | 0.002585754 | 0.002585754 |
| Sub18 | 5.3 | ductal | 0.241015599 | 0.241013942 | 0.241015599 |
| Sub18 | 5.3 | acinar | 0.068552743 | 0.068568287 | 0.068552743 |
| Sub19 | 4.5 | beta | 0.405271961 | 0.405588607 | 0.405271961 |
| Sub19 | 4.5 | acinar | 0.153343575 | 0.153329757 | 0.153343575 |
| Sub19 | 4.5 | gamma | 0.005157603 | 0.005156105 | 0.005157603 |
| Sub19 | 4.5 | delta | 0.015217274 | 0.015219433 | 0.015217274 |
| Sub19 | 4.5 | alpha | 0.183951205 | 0.183899082 | 0.183951205 |
| Sub19 | 4.5 | ductal | 0.237058382 | 0.236807016 | 0.237058382 |
| Sub21 | 4.3 | delta | 0.036630262 | 0.036636286 | 0.036630262 |
| Sub21 | 4.3 | beta | 0.674926802 | 0.674842582 | 0.674926802 |
| Sub21 | 4.3 | acinar | 0.015961302 | 0.015961186 | 0.015961302 |
| Sub21 | 4.3 | ductal | 0.08855 | 0.088605241 | 0.08855 |
| Sub21 | 4.3 | alpha | 0.183931634 | 0.183954704 | 0.183931634 |
| Sub21 | 4.3 | gamma | 0 | 0 | 0 |
| Sub23 | 5.4 | beta | 0.129388369 | 0.129299032 | 0.129388369 |
| Sub23 | 5.4 | alpha | 0.438808634 | 0.438989686 | 0.438808634 |
| Sub23 | 5.4 | acinar | 0.194361545 | 0.19437428 | 0.194361545 |
| Sub23 | 5.4 | gamma | 0.006360752 | 0.006359832 | 0.006360752 |
| Sub23 | 5.4 | delta | 0.026604052 | 0.026611858 | 0.026604052 |
| Sub23 | 5.4 | ductal | 0.204476648 | 0.204365312 | 0.204476648 |
| Sub24 | 5.5 | ductal | 0.347771349 | 0.347727451 | 0.347771349 |
| Sub24 | 5.5 | delta | 0.031086499 | 0.031094945 | 0.031086499 |
| Sub24 | 5.5 | alpha | 0.102959309 | 0.102909614 | 0.102959309 |
| Sub24 | 5.5 | gamma | 0.002972017 | 0.002971774 | 0.002972017 |
| Sub24 | 5.5 | beta | 0.419555611 | 0.419630032 | 0.419555611 |
| Sub24 | 5.5 | acinar | 0.095655216 | 0.095666183 | 0.095655216 |
| Sub25 | 6.2 | acinar | 0.376416367 | 0.211814391 | 0.25569629 |
| Sub25 | 6.2 | beta | 0.185226838 | 0.384596456 | 0.125098551 |
| Sub25 | 6.2 | alpha | 0.134331769 | 0.003931747 | 0.135084997 |
| Sub25 | 6.2 | delta | 0.007599718 | 0.009023637 | 0.01067841 |
| Sub25 | 6.2 | ductal | 0.292566451 | 0.388157839 | 0.470589987 |
| Sub25 | 6.2 | gamma | 0.003858856 | 0.00247593 | 0.002851765 |
| Sub26 | 5.8 | gamma | 0 | 0.000189176 | 0.000183659 |
| Sub26 | 5.8 | beta | 0.214088524 | 0.017961508 | 0.010279021 |
| Sub26 | 5.8 | alpha | 0.18178585 | 0.282660716 | 0.284054105 |
| Sub26 | 5.8 | delta | 0.001817068 | 0.003856559 | 0.003797841 |
| Sub26 | 5.8 | ductal | 0.115011011 | 0.311435849 | 0.333683741 |
| Sub26 | 5.8 | acinar | 0.487297548 | 0.383896192 | 0.368001633 |
| Sub27 | 6 | acinar | 0.199924418 | 0.168363996 | 0.150979238 |
| Sub27 | 6 | alpha | 0.185393345 | 0.166087905 | 0.216243697 |
| Sub27 | 6 | ductal | 0.181120179 | 0.204582398 | 0.260305103 |
| Sub27 | 6 | beta | 0.42388898 | 0.446466694 | 0.359425345 |
| Sub27 | 6 | delta | 0.009673078 | 0.014364667 | 0.012961203 |
| Sub27 | 6 | gamma | 0 | 0.000134339 | 8.54E-05 |
| Sub28 | 5.9 | gamma | 0 | 0 | 0 |
| Sub28 | 5.9 | acinar | 0.201005752 | 0.068839576 | 0.140540101 |
| Sub28 | 5.9 | ductal | 0.111278964 | 0.418907917 | 0.240489504 |
| Sub28 | 5.9 | alpha | 0.463271048 | 0.079492063 | 0.600778881 |
| Sub28 | 5.9 | delta | 0.00166783 | 0.003662285 | 0.004354254 |
| Sub28 | 5.9 | beta | 0.222776406 | 0.429098158 | 0.01383726 |
| Sub29 | 7 | alpha | 0.338175036 | 0.434489435 | 0.38438346 |
| Sub29 | 7 | beta | 0.489986058 | 0.372178303 | 0.374595713 |
| Sub29 | 7 | acinar | 0.062016166 | 0.038403294 | 0.042406748 |
| Sub29 | 7 | gamma | 0.002403772 | 0.001970108 | 0.002233007 |
| Sub29 | 7 | delta | 0.014745247 | 0.017139036 | 0.01913117 |
| Sub29 | 7 | ductal | 0.092673722 | 0.135819824 | 0.177249902 |
| Sub3 | 5.4 | alpha | 0.191348539 | 0.191483943 | 0.191348539 |
| Sub3 | 5.4 | beta | 0.352111748 | 0.35198963 | 0.352111748 |
| Sub3 | 5.4 | delta | 0.03203777 | 0.032046531 | 0.03203777 |
| Sub3 | 5.4 | acinar | 0.191436872 | 0.191466007 | 0.191436872 |
| Sub3 | 5.4 | gamma | 0.000311754 | 0.000311605 | 0.000311754 |
| Sub3 | 5.4 | ductal | 0.232753317 | 0.232702285 | 0.232753317 |
| Sub30 | 5.6 | alpha | 0.591235071 | 0.591509858 | 0.591235071 |
| Sub30 | 5.6 | gamma | 0.000629379 | 0.000629044 | 0.000629379 |
| Sub30 | 5.6 | beta | 0.230595241 | 0.230350074 | 0.230595241 |
| Sub30 | 5.6 | delta | 0.010143774 | 0.010148895 | 0.010143774 |
| Sub30 | 5.6 | ductal | 0.107418335 | 0.107364271 | 0.107418335 |
| Sub30 | 5.6 | acinar | 0.0599782 | 0.05999786 | 0.0599782 |
| Sub31 | 5.6 | gamma | 0.000328875 | 0.000328568 | 0.000328875 |
| Sub31 | 5.6 | acinar | 0.132238603 | 0.132152195 | 0.132238603 |
| Sub31 | 5.6 | alpha | 0.247218037 | 0.247153733 | 0.247218037 |
| Sub31 | 5.6 | ductal | 0.305928423 | 0.305671365 | 0.305928423 |
| Sub31 | 5.6 | beta | 0.309846405 | 0.310253938 | 0.309846405 |
| Sub31 | 5.6 | delta | 0.004439657 | 0.004440201 | 0.004439657 |
| Sub32 | 8.6 | delta | 0.006554859 | 0.009695744 | 0.009445818 |
| Sub32 | 8.6 | acinar | 0.29240522 | 0.221598986 | 0.212375035 |
| Sub32 | 8.6 | beta | 0.243337334 | 0.163578027 | 0.163177733 |
| Sub32 | 8.6 | alpha | 0.234211624 | 0.267165685 | 0.269028884 |
| Sub32 | 8.6 | ductal | 0.222093744 | 0.335950098 | 0.344008095 |
| Sub32 | 8.6 | gamma | 0.00139722 | 0.00201146 | 0.001964434 |
| Sub33 | 5.4 | acinar | 0.056192592 | 0.05620318 | 0.056192592 |
| Sub33 | 5.4 | alpha | 0.294756022 | 0.294902947 | 0.294756022 |
| Sub33 | 5.4 | beta | 0.472365501 | 0.47217966 | 0.472365501 |
| Sub33 | 5.4 | delta | 0.015315123 | 0.015323431 | 0.015315123 |
| Sub33 | 5.4 | ductal | 0.159352071 | 0.159372139 | 0.159352071 |
| Sub33 | 5.4 | gamma | 0.002018691 | 0.002018644 | 0.002018691 |
| Sub34 | 5.3 | ductal | 0.061149131 | 0.061070356 | 0.061149131 |
| Sub34 | 5.3 | delta | 0.024572882 | 0.024582593 | 0.024572882 |
| Sub34 | 5.3 | acinar | 0.164965387 | 0.164985262 | 0.164965387 |
| Sub34 | 5.3 | alpha | 0.475798068 | 0.475957563 | 0.475798068 |
| Sub34 | 5.3 | beta | 0.272623536 | 0.272513516 | 0.272623536 |
| Sub34 | 5.3 | gamma | 0.000890996 | 0.00089071 | 0.000890996 |
| Sub35 | 6.2 | acinar | 0.358191696 | 0.270115986 | 0.254191372 |
| Sub35 | 6.2 | alpha | 0.084638185 | 0.104904965 | 0.099886118 |
| Sub35 | 6.2 | beta | 0.357164515 | 0.297502116 | 0.334040941 |
| Sub35 | 6.2 | delta | 0.00328705 | 0.004706169 | 0.004345145 |
| Sub35 | 6.2 | ductal | 0.181672936 | 0.316801153 | 0.301944206 |
| Sub35 | 6.2 | gamma | 0.015045619 | 0.00596961 | 0.005592218 |
| Sub36 | 5 | gamma | 0.001995262 | 0.00199502 | 0.001995262 |
| Sub36 | 5 | ductal | 0.304964943 | 0.304801474 | 0.304964943 |
| Sub36 | 5 | acinar | 0.385395163 | 0.385447668 | 0.385395163 |
| Sub36 | 5 | alpha | 0.263674477 | 0.263742627 | 0.263674477 |
| Sub36 | 5 | beta | 0.026206686 | 0.02624563 | 0.026206686 |
| Sub36 | 5 | delta | 0.017763468 | 0.017767581 | 0.017763468 |
| Sub37 | 5.2 | acinar | 0.072524673 | 0.072520491 | 0.072524673 |
| Sub37 | 5.2 | alpha | 0.613058826 | 0.613293474 | 0.613058826 |
| Sub37 | 5.2 | beta | 0.147021965 | 0.147073926 | 0.147021965 |
| Sub37 | 5.2 | delta | 0.024351148 | 0.024355432 | 0.024351148 |
| Sub37 | 5.2 | ductal | 0.142538037 | 0.142251846 | 0.142538037 |
| Sub37 | 5.2 | gamma | 0.000505352 | 0.000504831 | 0.000505352 |
| Sub38 | 6 | ductal | 0.018409273 | 0.041054859 | 0.049148949 |
| Sub38 | 6 | delta | 0.022689228 | 0.029212921 | 0.028695564 |
| Sub38 | 6 | gamma | 0.017386745 | 0.010799368 | 0.01063274 |
| Sub38 | 6 | beta | 0.635767572 | 0.592294924 | 0.582657998 |
| Sub38 | 6 | acinar | 0.040976396 | 0.027221649 | 0.026342038 |
| Sub38 | 6 | alpha | 0.264770786 | 0.299416279 | 0.30252271 |
| Sub39 | 5.3 | beta | 0.540058433 | 0.540045724 | 0.540058433 |
| Sub39 | 5.3 | alpha | 0.095547037 | 0.095554561 | 0.095547037 |
| Sub39 | 5.3 | acinar | 0.149736337 | 0.149770525 | 0.149736337 |
| Sub39 | 5.3 | delta | 0.019186013 | 0.019194171 | 0.019186013 |
| Sub39 | 5.3 | ductal | 0.170664799 | 0.170630027 | 0.170664799 |
| Sub39 | 5.3 | gamma | 0.02480738 | 0.024804992 | 0.02480738 |
| Sub40 | 5.6 | acinar | 0.575456559 | 0.575425219 | 0.575456559 |
| Sub40 | 5.6 | alpha | 0.088577535 | 0.088562081 | 0.088577535 |
| Sub40 | 5.6 | beta | 0.081730022 | 0.081898197 | 0.081730022 |
| Sub40 | 5.6 | delta | 0.001240235 | 0.001240524 | 0.001240235 |
| Sub40 | 5.6 | ductal | 0.249664878 | 0.249543719 | 0.249664878 |
| Sub40 | 5.6 | gamma | 0.003330771 | 0.00333026 | 0.003330771 |
| Sub42 | 6.8 | delta | 0.010200241 | 0.014264278 | 0.016699739 |
| Sub42 | 6.8 | beta | 0.192155934 | 0.538642992 | 0.105980461 |
| Sub42 | 6.8 | acinar | 0.264769801 | 0.09670516 | 0.181348368 |
| Sub42 | 6.8 | alpha | 0.27707637 | 0.041436212 | 0.354178612 |
| Sub42 | 6.8 | ductal | 0.254117166 | 0.306560402 | 0.339089556 |
| Sub42 | 6.8 | gamma | 0.001680488 | 0.002390955 | 0.002703263 |
| Sub43 | 6.8 | acinar | 0.15985191 | 0.025903442 | 0.09846726 |
| Sub43 | 6.8 | alpha | 0.644083005 | 0.122472653 | 0.714094573 |
| Sub43 | 6.8 | beta | 0.089084365 | 0.49069823 | 0.002778588 |
| Sub43 | 6.8 | delta | 0.008458905 | 0.015912362 | 0.015313064 |
| Sub43 | 6.8 | ductal | 0.098429116 | 0.343950796 | 0.168585808 |
| Sub43 | 6.8 | gamma | 9.27E-05 | 0.001062517 | 0.000760707 |
| Sub44 | 5.5 | ductal | 0.314173127 | 0.314112025 | 0.314173127 |
| Sub44 | 5.5 | delta | 0.020007599 | 0.020011524 | 0.020007599 |
| Sub44 | 5.5 | acinar | 0.157824653 | 0.157838069 | 0.157824653 |
| Sub44 | 5.5 | alpha | 0.18029236 | 0.180326385 | 0.18029236 |
| Sub44 | 5.5 | beta | 0.327071009 | 0.327080939 | 0.327071009 |
| Sub44 | 5.5 | gamma | 0.000631251 | 0.000631057 | 0.000631251 |
| Sub45 | 5.3 | acinar | 0.028876241 | 0.028874282 | 0.028876241 |
| Sub45 | 5.3 | alpha | 0.528390318 | 0.528587549 | 0.528390318 |
| Sub45 | 5.3 | beta | 0.309320151 | 0.309196029 | 0.309320151 |
| Sub45 | 5.3 | delta | 0.022497896 | 0.022503306 | 0.022497896 |
| Sub45 | 5.3 | ductal | 0.104355291 | 0.104279922 | 0.104355291 |
| Sub45 | 5.3 | gamma | 0.006560104 | 0.006558911 | 0.006560104 |
| Sub46 | 8 | gamma | 0.001424592 | 0.002468418 | 0.002662671 |
| Sub46 | 8 | delta | 0.004025364 | 0.006479969 | 0.00694778 |
| Sub46 | 8 | ductal | 0.021738004 | 0.115090039 | 0.139490428 |
| Sub46 | 8 | beta | 0.159737625 | 0.121956454 | 0.130602374 |
| Sub46 | 8 | acinar | 0.498181197 | 0.343295633 | 0.355558291 |
| Sub46 | 8 | alpha | 0.314893218 | 0.410709488 | 0.364738456 |
| Sub47 | 4.6 | acinar | 0.213906966 | 0.213929181 | 0.213906966 |
| Sub47 | 4.6 | alpha | 0.158963542 | 0.158987246 | 0.158963542 |
| Sub47 | 4.6 | beta | 0.39734865 | 0.397319564 | 0.39734865 |
| Sub47 | 4.6 | delta | 0.016388751 | 0.016395762 | 0.016388751 |
| Sub47 | 4.6 | ductal | 0.207904749 | 0.207881053 | 0.207904749 |
| Sub47 | 4.6 | gamma | 0.005487342 | 0.005487196 | 0.005487342 |
| Sub48 | 6.2 | gamma | 0.045246147 | 0.025622611 | 0.025854777 |
| Sub48 | 6.2 | acinar | 0.071620302 | 0.048636735 | 0.048771859 |
| Sub48 | 6.2 | alpha | 0.257486307 | 0.279001038 | 0.258733713 |
| Sub48 | 6.2 | beta | 0.503271306 | 0.468697802 | 0.480492518 |
| Sub48 | 6.2 | delta | 0.013198668 | 0.017060198 | 0.016893479 |
| Sub48 | 6.2 | ductal | 0.10917727 | 0.160981616 | 0.169253654 |
| Sub50 | 5.6 | alpha | 0.382117712 | 0.382427088 | 0.382117712 |
| Sub50 | 5.6 | gamma | 0.000421963 | 0.000421577 | 0.000421963 |
| Sub50 | 5.6 | beta | 0.310120934 | 0.309856813 | 0.310120934 |
| Sub50 | 5.6 | delta | 0.006541787 | 0.006544249 | 0.006541787 |
| Sub50 | 5.6 | ductal | 0.213111731 | 0.21305009 | 0.213111731 |
| Sub50 | 5.6 | acinar | 0.087685873 | 0.087700184 | 0.087685873 |
| Sub51 | 6 | acinar | 0.126510871 | 0.085472986 | 0.083272842 |
| Sub51 | 6 | alpha | 0.216276001 | 0.235692173 | 0.224544098 |
| Sub51 | 6 | beta | 0.441299767 | 0.345693172 | 0.345091141 |
| Sub51 | 6 | delta | 0.007874589 | 0.010667246 | 0.010193996 |
| Sub51 | 6 | ductal | 0.20735528 | 0.321044718 | 0.335526284 |
| Sub51 | 6 | gamma | 0.000683493 | 0.001429705 | 0.001371639 |
| Sub52 | 5.5 | beta | 0.294277976 | 0.294179212 | 0.294277976 |
| Sub52 | 5.5 | alpha | 0.251987045 | 0.252084392 | 0.251987045 |
| Sub52 | 5.5 | acinar | 0.125432882 | 0.125483096 | 0.125432882 |
| Sub52 | 5.5 | delta | 0.052912257 | 0.052926808 | 0.052912257 |
| Sub52 | 5.5 | ductal | 0.273444468 | 0.273381316 | 0.273444468 |
| Sub52 | 5.5 | gamma | 0.001945372 | 0.001945176 | 0.001945372 |
| Sub53 | 6.2 | acinar | 0.512008008 | 0.372597266 | 0.35778244 |
| Sub53 | 6.2 | gamma | 0.006634696 | 0.004646346 | 0.004460035 |
| Sub53 | 6.2 | alpha | 0.12333842 | 0.143657688 | 0.138485579 |
| Sub53 | 6.2 | beta | 0.144322263 | 0.093472633 | 0.115957202 |
| Sub53 | 6.2 | delta | 0.003875805 | 0.005916148 | 0.005632421 |
| Sub53 | 6.2 | ductal | 0.209820807 | 0.379709919 | 0.377682323 |
| Sub54 | 6.4 | ductal | 0.120719463 | 0.194230294 | 0.195991444 |
| Sub54 | 6.4 | delta | 0.007717287 | 0.010588059 | 0.010292354 |
| Sub54 | 6.4 | alpha | 0.218968999 | 0.235904175 | 0.228262174 |
| Sub54 | 6.4 | beta | 0.38282314 | 0.371541429 | 0.38156867 |
| Sub54 | 6.4 | acinar | 0.268045585 | 0.185647195 | 0.181847683 |
| Sub54 | 6.4 | gamma | 0.001725525 | 0.002088848 | 0.002037675 |
| Sub55 | 7 | acinar | 0.391733883 | 0.264606871 | 0.277737302 |
| Sub55 | 7 | alpha | 0.095797662 | 0.106671159 | 0.093437408 |
| Sub55 | 7 | beta | 0.258558368 | 0.219343657 | 0.231723996 |
| Sub55 | 7 | delta | 0.008279184 | 0.011144929 | 0.011562224 |
| Sub55 | 7 | ductal | 0.222808455 | 0.385614903 | 0.372261699 |
| Sub55 | 7 | gamma | 0.022822448 | 0.012618481 | 0.013277371 |
| Sub56 | 10 | gamma | 0.001406129 | 0.002038317 | 0.001945406 |
| Sub56 | 10 | ductal | 0.321497425 | 0.52431354 | 0.544160821 |
| Sub56 | 10 | acinar | 0.534225963 | 0.355406068 | 0.345035661 |
| Sub56 | 10 | alpha | 0.09508138 | 0.115689387 | 0.106570397 |
| Sub56 | 10 | beta | 0.047279668 | 0.001466384 | 0.001255189 |
| Sub56 | 10 | delta | 0.000509434 | 0.001086303 | 0.001032525 |
| Sub57 | 5.1 | acinar | 0.091382552 | 0.091407227 | 0.091382552 |
| Sub57 | 5.1 | alpha | 0.222999747 | 0.222908683 | 0.222999747 |
| Sub57 | 5.1 | beta | 0.38626042 | 0.38634157 | 0.38626042 |
| Sub57 | 5.1 | delta | 0.005674793 | 0.005677287 | 0.005674793 |
| Sub57 | 5.1 | ductal | 0.293201384 | 0.293184242 | 0.293201384 |
| Sub57 | 5.1 | gamma | 0.000481104 | 0.00048099 | 0.000481104 |
| Sub58 | 5.3 | gamma | 0.002975229 | 0.002975036 | 0.002975229 |
| Sub58 | 5.3 | acinar | 0.038542654 | 0.038547151 | 0.038542654 |
| Sub58 | 5.3 | ductal | 0.280924341 | 0.280888206 | 0.280924341 |
| Sub58 | 5.3 | alpha | 0.110165513 | 0.11010169 | 0.110165513 |
| Sub58 | 5.3 | beta | 0.549552101 | 0.549642238 | 0.549552101 |
| Sub58 | 5.3 | delta | 0.017840162 | 0.017845678 | 0.017840162 |
| Sub59 | 5.4 | acinar | 0.217293076 | 0.217316662 | 0.217293076 |
| Sub59 | 5.4 | alpha | 0.115102963 | 0.114838127 | 0.115102963 |
| Sub59 | 5.4 | beta | 0.308928263 | 0.309213867 | 0.308928263 |
| Sub59 | 5.4 | delta | 0.013007745 | 0.01301152 | 0.013007745 |
| Sub59 | 5.4 | ductal | 0.33947132 | 0.339424116 | 0.33947132 |
| Sub59 | 5.4 | gamma | 0.006196633 | 0.006195708 | 0.006196633 |
| Sub6 | 5.5 | beta | 0.634553601 | 0.634578473 | 0.634553601 |
| Sub6 | 5.5 | acinar | 0.006650877 | 0.006652613 | 0.006650877 |
| Sub6 | 5.5 | alpha | 0.202667498 | 0.202705743 | 0.202667498 |
| Sub6 | 5.5 | delta | 0.003597401 | 0.0035997 | 0.003597401 |
| Sub6 | 5.5 | ductal | 0.151936441 | 0.151869371 | 0.151936441 |
| Sub6 | 5.5 | gamma | 0.000594182 | 0.000594101 | 0.000594182 |
| Sub60 | 5.3 | alpha | 0.147931573 | 0.147899987 | 0.147931573 |
| Sub60 | 5.3 | beta | 0.047626182 | 0.047771205 | 0.047626182 |
| Sub60 | 5.3 | delta | 0.014555024 | 0.014559323 | 0.014555024 |
| Sub60 | 5.3 | gamma | 0.000589182 | 0.000589055 | 0.000589182 |
| Sub60 | 5.3 | ductal | 0.500560186 | 0.500418783 | 0.500560186 |
| Sub60 | 5.3 | acinar | 0.288737853 | 0.288761647 | 0.288737853 |
| Sub61 | 6 | acinar | 0.609012573 | 0.478879195 | 0.47317977 |
| Sub61 | 6 | alpha | 0.194522806 | 0.241155071 | 0.167519606 |
| Sub61 | 6 | delta | 0.00753876 | 0.01092685 | 0.011416061 |
| Sub61 | 6 | ductal | 0.054940905 | 0.133199987 | 0.155296451 |
| Sub61 | 6 | gamma | 0.005659985 | 0.004392188 | 0.004622557 |
| Sub61 | 6 | beta | 0.128324971 | 0.131446709 | 0.187965554 |
| Sub62 | 5.8 | beta | 0.233839439 | 0.041455389 | 0.133297875 |
| Sub62 | 5.8 | delta | 0.006944438 | 0.012141515 | 0.011096091 |
| Sub62 | 5.8 | ductal | 0.05751854 | 0.085162689 | 0.122067451 |
| Sub62 | 5.8 | gamma | 0.000535553 | 0.002877839 | 0.002683062 |
| Sub62 | 5.8 | alpha | 0.67022409 | 0.838685968 | 0.708981514 |
| Sub62 | 5.8 | acinar | 0.030937939 | 0.0196766 | 0.021874007 |
| Sub63 | 5.4 | alpha | 0.700839863 | 0.701497056 | 0.700839863 |
| Sub63 | 5.4 | beta | 0.115295063 | 0.11479928 | 0.115295063 |
| Sub63 | 5.4 | acinar | 0.073740661 | 0.073765039 | 0.073740661 |
| Sub63 | 5.4 | ductal | 0.09829374 | 0.098100708 | 0.09829374 |
| Sub63 | 5.4 | gamma | 0 | 0 | 0 |
| Sub63 | 5.4 | delta | 0.011830673 | 0.011837917 | 0.011830673 |
| Sub64 | 5.7 | gamma | 0.000201379 | 0.001352743 | 0.001479286 |
| Sub64 | 5.7 | delta | 0.002237994 | 0.00308501 | 0.003600286 |
| Sub64 | 5.7 | ductal | 0.11905137 | 0.243139242 | 0.213452369 |
| Sub64 | 5.7 | acinar | 0.122376573 | 0.041223454 | 0.140397385 |
| Sub64 | 5.7 | alpha | 0.265786172 | 0.204611421 | 0.324235915 |
| Sub64 | 5.7 | beta | 0.490346513 | 0.50658813 | 0.316834759 |
| Sub65 | 6.2 | beta | 0.28327292 | 0.236288383 | 0.238649574 |
| Sub65 | 6.2 | delta | 0.005155645 | 0.007382097 | 0.007040602 |
| Sub65 | 6.2 | acinar | 0.10446891 | 0.077111816 | 0.07245227 |
| Sub65 | 6.2 | alpha | 0.15169423 | 0.182121532 | 0.174851398 |
| Sub65 | 6.2 | ductal | 0.303278793 | 0.42145659 | 0.434792008 |
| Sub65 | 6.2 | gamma | 0.152129502 | 0.075639582 | 0.072214148 |
| Sub66 | 5.2 | acinar | 0.052506036 | 0.052511281 | 0.052506036 |
| Sub66 | 5.2 | alpha | 0.052056624 | 0.052051843 | 0.052056624 |
| Sub66 | 5.2 | ductal | 0.360259535 | 0.360201784 | 0.360259535 |
| Sub66 | 5.2 | gamma | 0.010136199 | 0.010135738 | 0.010136199 |
| Sub66 | 5.2 | beta | 0.510606329 | 0.510658015 | 0.510606329 |
| Sub66 | 5.2 | delta | 0.014435277 | 0.014441339 | 0.014435277 |
| Sub67 | 5.7 | acinar | 0.373251162 | 0.083203382 | 0.257146892 |
| Sub67 | 5.7 | alpha | 0.122840836 | 0.014073431 | 0.116838497 |
| Sub67 | 5.7 | ductal | 0.118366977 | 0.176420173 | 0.248435553 |
| Sub67 | 5.7 | gamma | 0.001792904 | 0.001738845 | 0.001832345 |
| Sub67 | 5.7 | beta | 0.381420442 | 0.721526445 | 0.372296213 |
| Sub67 | 5.7 | delta | 0.002327678 | 0.003037724 | 0.003450499 |
| Sub68 | 5.2 | alpha | 0.251262805 | 0.251368655 | 0.251262805 |
| Sub68 | 5.2 | beta | 0.473769497 | 0.47369561 | 0.473769497 |
| Sub68 | 5.2 | delta | 0.022524123 | 0.022531864 | 0.022524123 |
| Sub68 | 5.2 | acinar | 0.085376213 | 0.085389342 | 0.085376213 |
| Sub68 | 5.2 | gamma | 0.004749417 | 0.004749008 | 0.004749417 |
| Sub68 | 5.2 | ductal | 0.162317945 | 0.162265521 | 0.162317945 |
| Sub69 | 5.4 | alpha | 0.317656261 | 0.317762497 | 0.317656261 |
| Sub69 | 5.4 | beta | 0.450579359 | 0.450512845 | 0.450579359 |
| Sub69 | 5.4 | acinar | 0.051582765 | 0.051592252 | 0.051582765 |
| Sub69 | 5.4 | gamma | 0.000782118 | 0.000781931 | 0.000782118 |
| Sub69 | 5.4 | delta | 0.031612116 | 0.031624472 | 0.031612116 |
| Sub69 | 5.4 | ductal | 0.14778738 | 0.147726003 | 0.14778738 |
| Sub70 | 5.6 | acinar | 0.188230652 | 0.188241677 | 0.188230652 |
| Sub70 | 5.6 | beta | 0.107330828 | 0.107532053 | 0.107330828 |
| Sub70 | 5.6 | delta | 0.005910839 | 0.005912851 | 0.005910839 |
| Sub70 | 5.6 | ductal | 0.313415513 | 0.313260398 | 0.313415513 |
| Sub70 | 5.6 | alpha | 0.383240564 | 0.383181857 | 0.383240564 |
| Sub70 | 5.6 | gamma | 0.001871603 | 0.001871164 | 0.001871603 |
| Sub71 | 5.6 | alpha | 0.514481672 | 0.514794037 | 0.514481672 |
| Sub71 | 5.6 | beta | 0.004038719 | 0.0037446 | 0.004038719 |
| Sub71 | 5.6 | delta | 0.010142019 | 0.010148165 | 0.010142019 |
| Sub71 | 5.6 | gamma | 0.000165315 | 0.000164978 | 0.000165315 |
| Sub71 | 5.6 | ductal | 0.328954869 | 0.328850048 | 0.328954869 |
| Sub71 | 5.6 | acinar | 0.142217406 | 0.142298171 | 0.142217406 |
| Sub72 | 5.8 | acinar | 0.026856668 | 0.006426236 | 0.017311059 |
| Sub72 | 5.8 | alpha | 0.197732273 | 0.042527524 | 0.154625299 |
| Sub72 | 5.8 | beta | 0.693185529 | 0.809175715 | 0.70103781 |
| Sub72 | 5.8 | delta | 0.005544353 | 0.007190704 | 0.007008972 |
| Sub72 | 5.8 | ductal | 0.073554482 | 0.131976927 | 0.117425329 |
| Sub72 | 5.8 | gamma | 0.003126694 | 0.002702892 | 0.002591532 |
| Sub73 | 5.7 | beta | 0.118827427 | 0.005679371 | 0.020277493 |
| Sub73 | 5.7 | delta | 0.004172143 | 0.008447585 | 0.007857145 |
| Sub73 | 5.7 | ductal | 0.147439929 | 0.249421877 | 0.232520765 |
| Sub73 | 5.7 | gamma | 0 | 0.002123332 | 0.001992364 |
| Sub73 | 5.7 | acinar | 0.144950013 | 0.048299542 | 0.083573515 |
| Sub73 | 5.7 | alpha | 0.584610488 | 0.686028293 | 0.653778718 |
| Sub74 | 5.6 | beta | 0.000899819 | 0.000900277 | 0.000899819 |
| Sub74 | 5.6 | delta | 0.002576521 | 0.002577621 | 0.002576521 |
| Sub74 | 5.6 | acinar | 0.299277764 | 0.299326736 | 0.299277764 |
| Sub74 | 5.6 | alpha | 0.207665494 | 0.207687229 | 0.207665494 |
| Sub74 | 5.6 | gamma | 0.002123859 | 0.002123725 | 0.002123859 |
| Sub74 | 5.6 | ductal | 0.487456543 | 0.487384411 | 0.487456543 |
| Sub75 | 5.7 | gamma | 0.003792779 | 0.00285618 | 0.002766144 |
| Sub75 | 5.7 | alpha | 0.143043897 | 0.144071222 | 0.139448185 |
| Sub75 | 5.7 | ductal | 0.394632107 | 0.501623099 | 0.521976888 |
| Sub75 | 5.7 | acinar | 0.049427114 | 0.035350294 | 0.032365393 |
| Sub75 | 5.7 | beta | 0.405569648 | 0.311125764 | 0.298643945 |
| Sub75 | 5.7 | delta | 0.003534454 | 0.004973442 | 0.004799444 |
| Sub76 | 6.1 | acinar | 0.07250323 | 0.056879213 | 0.051414856 |
| Sub76 | 6.1 | delta | 0.026027744 | 0.036998596 | 0.037081818 |
| Sub76 | 6.1 | ductal | 0.046351106 | 0.080272326 | 0.098904775 |
| Sub76 | 6.1 | alpha | 0.375569262 | 0.417279985 | 0.420816152 |
| Sub76 | 6.1 | beta | 0.476558063 | 0.405023512 | 0.388221505 |
| Sub76 | 6.1 | gamma | 0.002990595 | 0.003546367 | 0.003560894 |
| Sub77 | 6.1 | gamma | 0.000473006 | 0.001324166 | 0.001304574 |
| Sub77 | 6.1 | alpha | 0.225532155 | 0.245249325 | 0.243003477 |
| Sub77 | 6.1 | beta | 0.325674563 | 0.2473515 | 0.220230963 |
| Sub77 | 6.1 | acinar | 0.144218029 | 0.100361296 | 0.094780123 |
| Sub77 | 6.1 | delta | 0.004827797 | 0.006834293 | 0.006713028 |
| Sub77 | 6.1 | ductal | 0.29927445 | 0.39887942 | 0.433967835 |
| Sub78 | 7.3 | acinar | 0.024046767 | 0.016403459 | 0.016716964 |
| Sub78 | 7.3 | alpha | 0.469810011 | 0.50610583 | 0.482872903 |
| Sub78 | 7.3 | ductal | 0.100418971 | 0.139653253 | 0.162503856 |
| Sub78 | 7.3 | gamma | 0 | 0.00185123 | 0.001912199 |
| Sub78 | 7.3 | beta | 0.394369051 | 0.319503458 | 0.319178227 |
| Sub78 | 7.3 | delta | 0.0113552 | 0.01648277 | 0.016815852 |
| Sub8 | 4.3 | acinar | 0.087810189 | 0.087822268 | 0.087810189 |
| Sub8 | 4.3 | gamma | 0.004738344 | 0.004737928 | 0.004738344 |
| Sub8 | 4.3 | alpha | 0.140230612 | 0.140221336 | 0.140230612 |
| Sub8 | 4.3 | beta | 0.642634933 | 0.642574562 | 0.642634933 |
| Sub8 | 4.3 | delta | 0.018646574 | 0.018653324 | 0.018646574 |
| Sub8 | 4.3 | ductal | 0.105939348 | 0.105990581 | 0.105939348 |
| Sub80 | 5.7 | acinar | 0.400623108 | 0.303450564 | 0.296871406 |
| Sub80 | 5.7 | alpha | 0.219700191 | 0.284217988 | 0.278987466 |
| Sub80 | 5.7 | beta | 0.112870578 | 0.001270917 | 0.001187773 |
| Sub80 | 5.7 | delta | 0.002282607 | 0.004226474 | 0.004167021 |
| Sub80 | 5.7 | ductal | 0.264523515 | 0.406756549 | 0.418745621 |
| Sub80 | 5.7 | gamma | 0 | 7.75E-05 | 4.07E-05 |
| Sub81 | 5.5 | acinar | 0.05344482 | 0.053441488 | 0.05344482 |
| Sub81 | 5.5 | alpha | 0.434979177 | 0.43501748 | 0.434979177 |
| Sub81 | 5.5 | beta | 0.296427441 | 0.296511826 | 0.296427441 |
| Sub81 | 5.5 | delta | 0.011765075 | 0.011767713 | 0.011765075 |
| Sub81 | 5.5 | ductal | 0.202650615 | 0.202529082 | 0.202650615 |
| Sub81 | 5.5 | gamma | 0.000732871 | 0.000732411 | 0.000732871 |
| Sub82 | 7.2 | acinar | 0.245599644 | 0.272327752 | 0.229479528 |
| Sub82 | 7.2 | alpha | 0.061235391 | 0.064161536 | 0.059390431 |
| Sub82 | 7.2 | beta | 0.251076396 | 0.184187331 | 0.173937498 |
| Sub82 | 7.2 | delta | 0.012494488 | 0.017074568 | 0.015232063 |
| Sub82 | 7.2 | ductal | 0.385728332 | 0.45148338 | 0.512463998 |
| Sub82 | 7.2 | gamma | 0.043865748 | 0.010765434 | 0.009496483 |
| Sub83 | 6.4 | acinar | 0.385653876 | 0.321455745 | 0.256829165 |
| Sub83 | 6.4 | alpha | 0.136748698 | 0.208186295 | 0.14549221 |
| Sub83 | 6.4 | beta | 0.175141975 | 0.009938688 | 0.155069165 |
| Sub83 | 6.4 | delta | 0.001156439 | 0.002459252 | 0.001866605 |
| Sub83 | 6.4 | ductal | 0.301299012 | 0.45795281 | 0.440742856 |
| Sub83 | 6.4 | gamma | 0 | 7.21E-06 | 0 |
| Sub84 | 6.7 | acinar | 0.129045918 | 0.093209615 | 0.087953548 |
| Sub84 | 6.7 | alpha | 0.141682901 | 0.147740073 | 0.138019081 |
| Sub84 | 6.7 | beta | 0.462321829 | 0.365027942 | 0.372658965 |
| Sub84 | 6.7 | delta | 0.003009935 | 0.004014382 | 0.004208383 |
| Sub84 | 6.7 | ductal | 0.262743515 | 0.388682605 | 0.395771617 |
| Sub84 | 6.7 | gamma | 0.001195902 | 0.001325383 | 0.001388405 |
| Sub85 | 5.9 | acinar | 0.129153287 | 0.08606655 | 0.075481708 |
| Sub85 | 5.9 | alpha | 0.331077138 | 0.29546224 | 0.441381073 |
| Sub85 | 5.9 | beta | 0.314734467 | 0.283231583 | 0.145967858 |
| Sub85 | 5.9 | delta | 0.004419561 | 0.006924054 | 0.007846678 |
| Sub85 | 5.9 | ductal | 0.220430358 | 0.326784215 | 0.3276527 |
| Sub85 | 5.9 | gamma | 0.000185189 | 0.001531358 | 0.001669983 |
| Sub86 | 5.6 | acinar | 0.115098686 | 0.115098465 | 0.115098686 |
| Sub86 | 5.6 | alpha | 0.433115882 | 0.433189383 | 0.433115882 |
| Sub86 | 5.6 | beta | 0.220554306 | 0.220581899 | 0.220554306 |
| Sub86 | 5.6 | delta | 0.007894122 | 0.007896231 | 0.007894122 |
| Sub86 | 5.6 | ductal | 0.22325668 | 0.223154033 | 0.22325668 |
| Sub86 | 5.6 | gamma | 8.03E-05 | 8.00E-05 | 8.03E-05 |
| Sub87 | 6 | acinar | 0.067863689 | 0.043313474 | 0.044495993 |
| Sub87 | 6 | alpha | 0.181744642 | 0.193498054 | 0.194478639 |
| Sub87 | 6 | beta | 0.349057316 | 0.239347096 | 0.248503432 |
| Sub87 | 6 | delta | 0.006216603 | 0.007824538 | 0.007938742 |
| Sub87 | 6 | ductal | 0.368526801 | 0.506457095 | 0.494902672 |
| Sub87 | 6 | gamma | 0.026590949 | 0.009559743 | 0.009680523 |
| Sub88 | 5.6 | acinar | 0.011996879 | 0.012002451 | 0.011996879 |
| Sub88 | 5.6 | alpha | 0.574283191 | 0.574783844 | 0.574283191 |
| Sub88 | 5.6 | beta | 0.333517236 | 0.333031412 | 0.333517236 |
| Sub88 | 5.6 | delta | 0.018392635 | 0.0184014 | 0.018392635 |
| Sub88 | 5.6 | ductal | 0.045142392 | 0.045113803 | 0.045142392 |
| Sub88 | 5.6 | gamma | 0.016667666 | 0.01666709 | 0.016667666 |
| Sub89 | 5.3 | acinar | 0.010100628 | 0.010102486 | 0.010100628 |
| Sub89 | 5.3 | alpha | 0.367896256 | 0.368076801 | 0.367896256 |
| Sub89 | 5.3 | beta | 0.499555638 | 0.499364745 | 0.499555638 |
| Sub89 | 5.3 | delta | 0.014865387 | 0.014873174 | 0.014865387 |
| Sub89 | 5.3 | ductal | 0.103485495 | 0.103486075 | 0.103485495 |
| Sub89 | 5.3 | gamma | 0.004096596 | 0.00409672 | 0.004096596 |
